# Single-cell Bayesian deconvolution

**DOI:** 10.1101/2022.02.13.480250

**Authors:** Gabriel Torregrosa, David Oriola, Vikas Trivedi, Jordi Garcia-Ojalvo

## Abstract

Individual cells exhibit substantial heterogeneity in protein abundance and activity, which is frequently reflected in broad distributions of fluorescently labeled reporters. Since all cellular components are intrinsically fluorescent to some extent, the observed distributions contain background noise that masks the natural heterogeneity of cellular populations. This limits our ability to characterize cell-fate decision processes that are key for development, immune response, tissue homeostasis, and many other biological functions. It is therefore important to separate the contributions from signal and noise in single-cell measurements. Addressing this issue rigorously requires deconvolving the noise distribution from the signal, but approaches in that direction are still limited. Here we present a non-parametric Bayesian formalism that performs such a deconvolution efficiently on multidimensional measurements, in a way that allows us to estimate confidence intervals. We use the approach to study the expression of the mesodermal transcription factor Brachyury in mouse embryonic stem cells undergoing differentiation.

## INTRODUCTION

The inherent stochasticity of biological processes leads to substantial heterogeneity even among genetically identical cells in the same environment^1–3^. The degree to which this heterogeneity affects, or even dictates, cellular decision-making in most situations is still an open question. This issue is of paramount importance in processes such as mammalian development, where hundreds (if not thousands^4^) of distinct cell types (cell states) emerge from a small number of identical undifferentiated cells^5–7^. Identifying the molecular mechanisms underlying these cell-fate decision programs and their interplay with cellular heterogeneity^8,9^ requires a rigorous quantification of cellular states across large numbers of cells.

Flow cytometry enables monitoring the distributions of abundances and activities of selected proteins for thousands of cells at a time, using fluorescently labeled markers. Cells, however, have a non-negligible amount of autofluorescence in the emission spectrum of most fluorescent probes (500 to 700 nm). This leads to a background noise that must be subtracted from the total signal emitted by fluorescently labeled cells^10^, in order to adequately relate the signal distribution provided by the cytometer to the mechanisms regulating the expression and/or activity of the protein of interest. Several standard methods exist for addressing this issue on a cellby-cell basis. One can use, for instance, additional fluorescence channels outside the emission spectrum of the fluorophores serving as regressor variables^11,12^. Alternatively, a second laser system can provide an independent measurement of both signal and autofluorescence^13^. Beyond issues of cost or accessibility, the effectivity of such solutions is limited, because there is no guarantee that autofluorescence from another channel, or from another excitation source, is a good proxy for autofluorescence in our channel of interest.

Commonly, rather than dedicating measurement resources to assess autofluorescence, control measurements of unlabeled cells are used to set a baseline of the signal coming from the naturally present autofluorescent components in the cell. This procedure, however, does not lead to a quantitative determination of the distribution of the signal coming *exclusively* from the fluorescent probe. Such quantitative assessment would require deconvolving the fluorescence distribution obtained in labeled cells from the one produced by unlabeled cells. Here, we propose a non-parametric Bayesian approach to this deconvolution problem, applicable to multichannel measurements. The method is robust and efficient, requiring cell numbers not larger than those typically considered in standard flow cytometry runs, and gives natural confidence intervals of the target distributions, which makes it attractive for a variety of applications.

There is an extensive statistics literature addressing the additive deconvolution problem^14^. A common set of deconvolution methods are kernel-based approaches, such as those relying on Fourier transforms^15–21^, which use the fact that in Fourier space, a deconvolution is simply the product of two functions. Two problems arise from such methods that limit their applicability in practical cases. First, Fourier transforms (and other methods that use orthogonal local bases, such as wavelets^22^) are not positive defined. Consequently, kernel-based methods lead to deconvolved pseudo-distributions with artificial features, which are hardly interpretable for practical applications. Second, these methods usually lead to point estimates, and therefore do not provide native confidence intervals (i.e. without applying additional statistical approximations) that allow us to assess the quality of the inferred target distribution.

A second class of deconvolution approaches are likelihood-based methods^23^, which estimate the unknown target distribution using maximum likelihood approaches. As in the case of kernel-based methods, these approaches provide us with point estimates, and usually assume exact knowledge of the noise distribution. Finally, a third class of methods involve Bayesian inference^24–26^, which does not require complete knowledge of the noise distribution and naturally provides confidence intervals of the estimates obtained. So far, however, these Bayesian methods have been applied to repeated measurements of the same individual entities that are being monitored (in our case, cells), which is not a realistic possibility in standard flow cytometry. Our nonparametric Bayesian approach does not require repeated measurements and retains all the above-mentioned advantages of Bayesian methods. We have implemented the procedure in a Julia package available in GitHub (https://github.com/dsb-lab/scBayesDeconv.jl).

The work is structured as follows. First, we introduce the mathematical description of the additive convolution problem in the context of flow cytometry data, and explain our proposed deconvolution method based on non-parametric Bayesian models. Second, we validate our method using synthetic datasets with known target distributions, and compare its results to other existing methods. Third, we further test our method in real flowcytometry data of mouse embryonic stem cells undergoing differentiation. We test the dataset in two conditions. Initially, we treat our cells with a low concentration of a fluorescent dye (which masks the real flow-cytometry signal and acts as an external noise), and validate our method by deconvolving the noise coming from the dye and comparing it to the control case where the dye was not added. This offers evidence of the robustness of the method in real applications while having a ground truth. Additionally, we perform the deconvolution using several channels from the dataset, to show the effectiveness of the method in multichannel measurements, and comment on the complementarity of our approach to other processing steps common in analysis pipelines of flow cytometry data. We conclude by discussing the limitations of the method and possible ways to improve it in future work.

## RESULTS

### Theoretical definition of the problem

Consider a population of cells containing fluorescent markers that label the abundances or activities (e.g. phosphorylation states) of certain proteins of interest. Flow cytometry measurements provide us with the distribution *pc*(*C*) of total fluorescence signal *C* emitted by each individual cell in the population, as measured by the flow cytometer detectors. These signals have two components: the fluorescence *T* emitted exclusively by the target fluorophores that report on the proteins of interest, and the autofluorescence *ξ* emitted by cellular components *other* than our fluorescent labels:

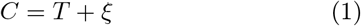

If these two components are independent of one another, the distribution *pc*(*C*) of the measured signal takes the form of a *convolution* of the distributions of *T* and *ξ*:

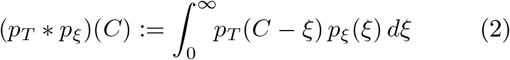

Similarly to *pc*, the distribution *pξ* of the autofluorescence *ξ* can be measured in the flow cytometer by using unlabeled cells that are otherwise identical to the labeled ones. On the other hand, the probability distribution of *T, pT*, cannot be measured directly. Our goal is to extract (deconvolve) the distribution *pT* from the measured distributions *pc* and *pξ*, considering that we only have a finite set of samples (cells) of *C* and *ξ*.

In what follows, we first introduce the way in which we describe the distributions involved in the problem. Next we define the posterior distribution given by the model and the data, and finally we discuss the methods used to explore the parameter space for the deconvolution problem.

#### Mixture model

The vast majority of real datasets (including those generated in flow cytometry) result from complex combinations of variables that cannot be explained in general with simple distributions. In order to adapt flexibly to such conditions, we use mixtures of probability distributions as our basis set. Any function can be arbitrarily well approximated using an adequate choice of basis functions, provided enough components are included in the mixture. We can describe both the target and the noise distributions by independent mixtures of *K* components:

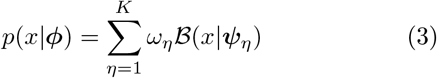

where *x* is multivariate in the case of multichannel measurements. *ωη* denotes the weight of each base *B*(*x*|***ψ****η*) in the mixture, and ***ψ****η* represents its parameters (the means and covariance matrices in the case of multivariate normal distributions)^27^. The sets of *ωη* and ***ψ****η* are in turn represented by the vector ***φ*** in what follows. Using these basis functions, the distribution of the observed signal *C* can be described as a superposition of all the convolutions between them:

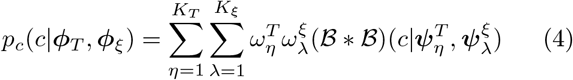

where 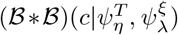 represents the convolution of two basis distributions with parameters 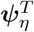 and 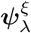, respectively, and *KT* and *Kξ* denote the number of bases used in each of the two mixtures.

The choice of basis functions to describe our data is crucial. We propose to use multivariate normal distributions to describe the data. Normal distributions with unknown mean and covariance have been shown to be flexible enough to represent datasets with high quality, requiring less components than more rigid methods. They have the additional advantage that its convolution turns out to be another normal distribution with modified parameters^28^. This last property is especially interesting for deriving exact analytic results for the Bayesian sampling process.

#### Posterior distribution and likelihood

Our goal is to extract, starting from samples of the distributions of total signal *C* and noise *ξ*, the parameters of the distribution of the target signal *T*. Defining the problem in terms of probability distributions leads very naturally to work with Bayesian methods. According to Bayes’ rule, the posterior distribution that represents the probability of the parameters given the data is

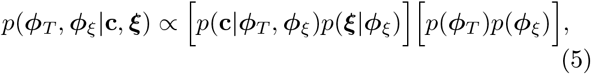

where **c** = {*ci* : *i* = 1 … *Nc*} and ***ξ*** = {*ξi* : *i* = 1 … *Nξ*} are the sets of observed samples of the total signal and the noise, respectively, with *Nc* and *Nξ* representing the number of samples in each case.

The first bracket on the right-hand side of Eq. (5) is the likelihood function, which corresponds to the probability of the data given the parameters. Under the assumption of independent and identically distributed (iid) observations, this function is given by

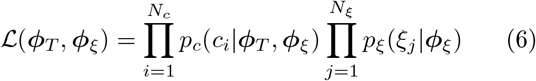

As can be seen, the information about the noise parameters is contained in both datasets, while the information about the target parameters only appears in the convolved data.

As shown in Eq. (5) above, the posterior distribution can be estimated by multiplying the likelihood by the prior distribution of parameter values. Since the datasets required for a good deconvolution are large, the impact of the prior distribution should be negligible. Therefore, the posterior landscape is effectively described by (6), and the prior distribution is only present in our approach for formal and computational reasons. In any case, since no prior information exists on the parameters that describe the noise and target mixture components, we impose vague priors over the plausible set of parameters (see Supplementary Sec. S4).

#### Sampling

The posterior distribution is a complex multimodal (multi-peaked) object containing all the configurations of the target and noise distributions that are consistent with the observed data. A correct and efficient exploration of this distribution is essential to set bounds on the candidate models that explain the data. The Bayesian model, as stated in Eq. (5), has no direct closed form. Sampling from this model as stated would require the introduction of complex sampling algorithms such as Markov Chain Monte Carlo (MCMC)^28^ and nested sampling^29^. These sampling processes are computationally very expensive and scale linearly with the sample size, which in the case of flow cytometry datasets is of the orders of tens to hundreds of thousands samples per dataset.

In order to overcome these shortcomings, we propose two approximations to the posterior distribution. The first one decouples the inference of the noise and target distributions parameters as two independent problems (see Supplementary Section S2.1). The parameters of the noise distribution can be fitted using well-known mixtures of normal distributions, with the target being described by a convolved mixture model. The second approximation simplifies the sampling of the convolved mixture model to enable sampling using standard Gibbs sampling algorithms (see Supplementary Section S4.2). The combination of both approximations allows the posterior to be sampled using Gibbs sampling, which is an efficient MCMC method of sampling from the posterior. In addition to these approximations, we offer two alternative Bayesian mixtures, namely finite or infinite normal mixture models^30^. The latter approach has the added advantage over the former of not requiring fixing the number of basis functions to describe the data. However, for most practical purposes, finite normal mixture models are able to fit the data with high accuracy and in a more efficient manner regarding how the corresponding algorithms work.

#### Analysis pipeline

Given the concepts and tools described above, the analysis of the data is performed as follows (Fig. 1). First, the distributions of the target and autofluorescence signals are assumed to be described by mixtures (top left panel in Fig. 1), as defined by Eqs. (3) and (4). The data measured experimentally correspond to the autofluorescence signal (noise) and the total signal of the labeled cells (which includes the autofluorescence), as shown in the top right panel of Fig. 1. The mixture assumption and the observed data samples allow us to construct the posterior distribution (Eq. (5) and middle panel in Fig. 1) from the likelihood function defined by Eq. (6) and the prior distributions discussed in the Supplementary Sec. S4. The posterior distribution is a function of the model parameters (weights of the mixtures and parameters of the basis functions). We next explore (sample) the posterior distribution in parameter space using the Gibbs sampling approximation (bottom left panels in Fig. 1). This algorithm provides us with a representative sampling of the parameters of the target distribution, which allows us to compute an average of this distribution and its confidence interval (bottom right panel in the figure).

**FIG. 1.**
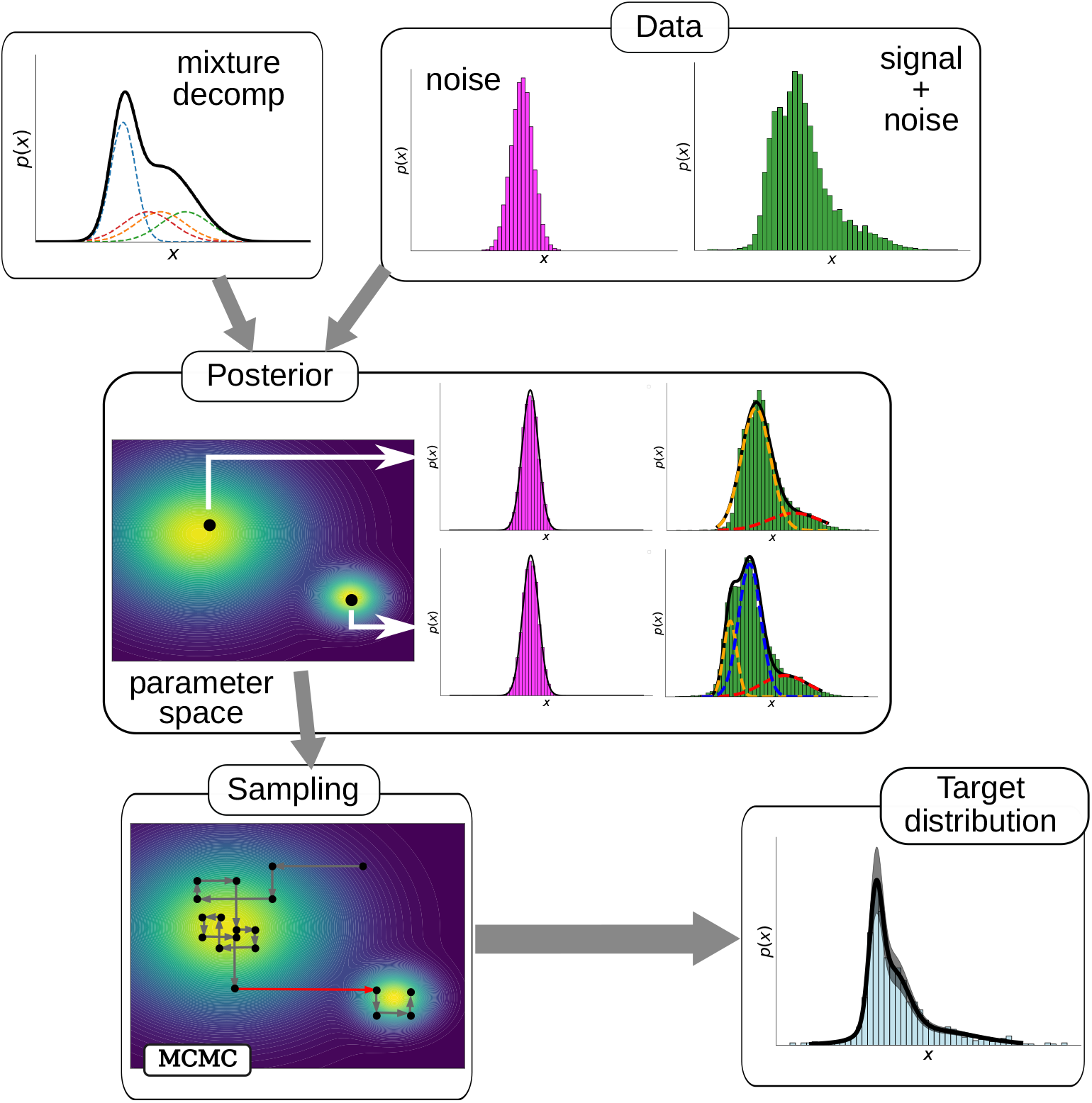
Scheme of the deconvolution process. The signal and noise mixture distributions, together with the observed data (top row), define the posterior distribution over the parameter space of the mixture, Eq. (6) (middle row). This distribution can present multiple peaks, sometimes degenerate with respect to basis label exchange, each corresponding to a different mixture description of the observed and target distributions. The red arrow in the Gibbs Markov Chain Monte Carlo sampling plot (bottom left) represents an unlikely jump between two peaks separated by a relatively wide probability valley.

### Application to synthetic data

First, we benchmark the ability of our method to recover the target distribution by using collections of synthetic datasets. In those datasets, the target and noise distributions are known, so we can compare the result of the deconvolution against a ground truth. To do this, we applied our method to the synthetic data described in Sec.. To prove the robustness of the implementation, we ran the test using four components, both for the noise and target distributions. We also avoided checking the full convergence of the algorithm manually. We ran the algorithm in this suboptimal conditions to avoid having to fine tune the specific parameters, which could lead to positive bias favoring our method in comparison with FFT-based approaches. We contrasted the results of our method with those of a specific FFT approach that does not require knowledge of the autofluorescence distribution^19^. Figure 2 shows a typical instance of the deconvolution performance of the two methods. Panel (a) shows in green the total (convolved) signal mimicking the output of a flow cytometry experiment. In this synthetic case, the signal is obtained by forward convolving a target distribution with the characteristics given above, shown in light blue in panels (b) and (c), with a noise (autofluorescence) distribution, shown in magenta in the inset of panel (a). The goal in this case is to recover (deconvolve) the ground-truth target distribution (panels b and c in Fig. 2) from the total and noise distributions (panel a). Figures 2(b,c) show that our Bayesian deconvolution method recovers the target distribution well. In comparison, the FFT method leads to oscillatory components in the deconvolution, and correspondingly to artefactual negative values in the probability distribution. Additionally, the Bayesian deconvolution method naturally provides a confidence i nterval, s hown by t he d ifferent samples (red lines) in panel (b) of Fig. 2.

**FIG. 2.**
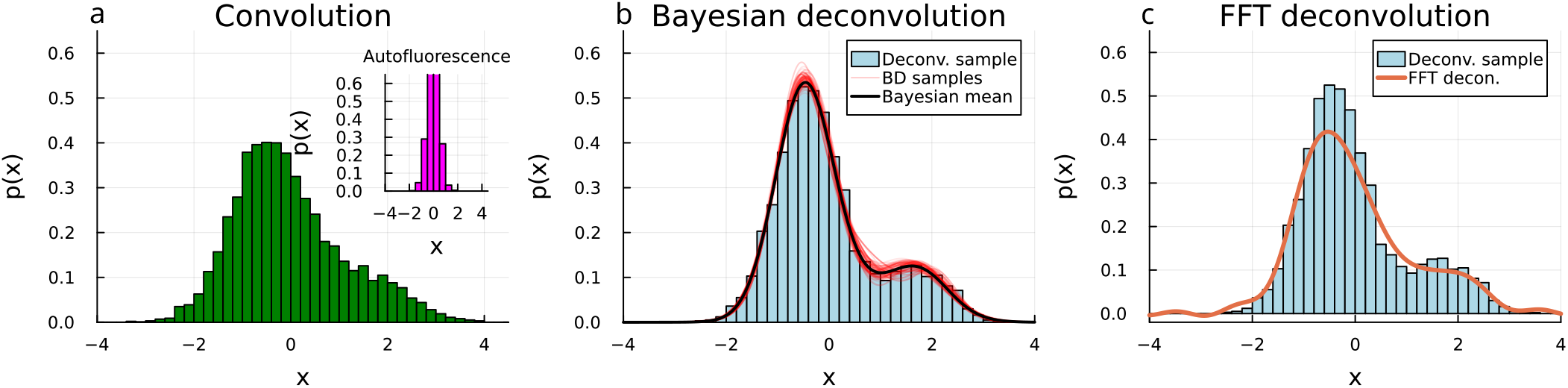
Deconvolution instance for a target bimodal distribution (light blue in panels b and c) corrupted by a normally distributed noise with a SNS = 2 (magenta distribution in the inset of panel a), applying the Bayesian Deconvolution method (panel b) and the FFT method (panel c). The total distribution from which the noise is deconvolved is shown in green in panel (a). The red lines in panel (b) depict samples of the Bayes fitting process. In this case, the noise distribution is a single normal function with mean *µ*^*ξ*^ = 0 and standard deviation *σ*^*ξ*^ = 0.5, and the target is a mixture of two normal functions with means 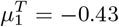 and 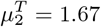, standard deviations 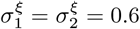, and weights 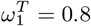 and 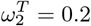.

For benchmarking purposes, we compared the deconvolved distribution with the real target distribution, which is known in this case, using the mean integrated overlap (MIO), as defined in Supplementary Sec. S 6. We preferred this measure to the mean integrated squared error (MISE), which is commonly used in the deconvolution literature for theoretical reasons^16,19,20,22^, since the MIO measure is easier to interpret, as it corresponds directly to the absolute overlap of two probability distributions. We also avoid more common measures of distribution dissimilarity such as the Kolmogorov-Smirnov test, since such methods would underestimate the ability of FFT-based methods to converge to the ground-truth deconvolution, given that they lead to artifacts in the resulting distributions, as shown above.

Figure 3 compares the deconvolution efficiency of the FFT-based method and our single-cell Bayesian deconvolution approach, in terms of the MIO measure that quantifies the similarity between the convolved distribution and the real one. According to its definition (see Supplementary Sec. S6), a MIO value of 1 corresponds to a perfect overlap, while the measure is 0 when two distributions do not overlap at all. As can be seen in Fig. 3, the single-cell Bayesian Deconvolution method outperforms the Fourier-based method in almost all the cases considered, the difference being more substantial for high levels of noise (small SNR, circles). In particular, the overlap is never below 0.7 for the Bayesian methods, while it can reach values near 0 in the FFT case depending on the type of distributions involved, particularly for low sampling numbers.

**FIG. 3.**
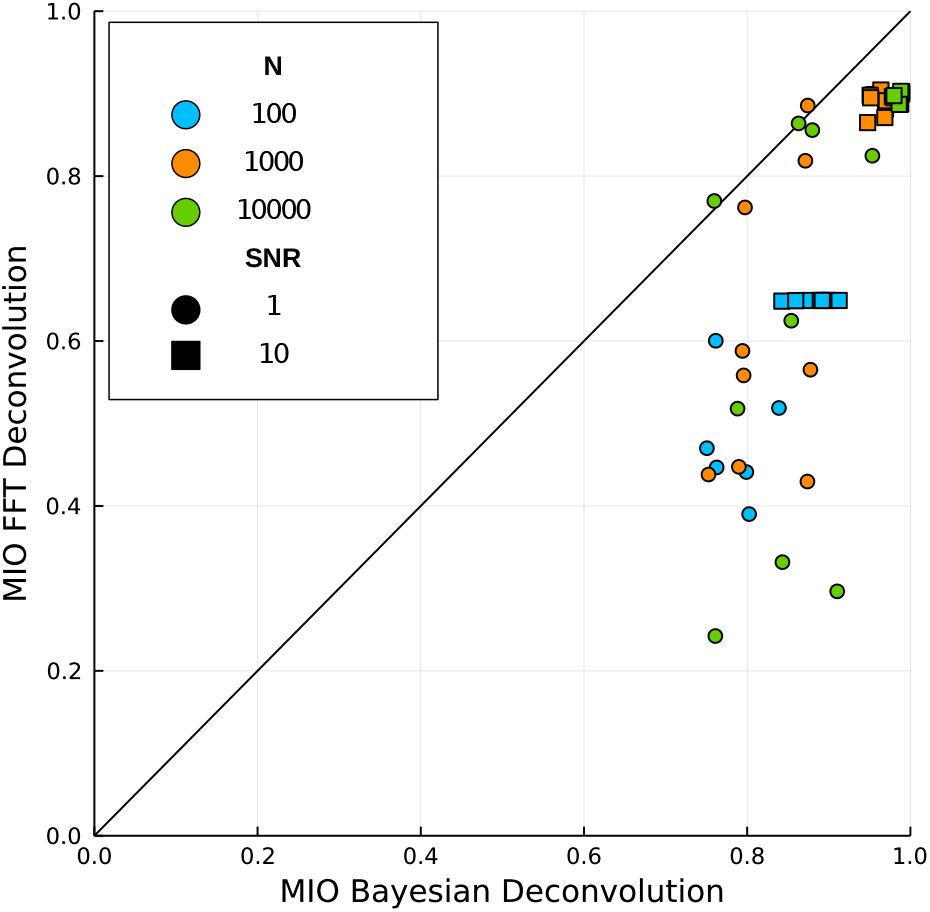
Similarity between the deconvolved and groundtruth target distributions as expressed by the Mean Integrated Overlap (MIO) for the two deconvolution methods (x and y axis), at different sampling sizes for the synthetic datasets described in Sec., with SNR = 1 (circles) and SNR = 10 (squares).

Additionally, it is worth noting that while the singlecell Bayesian deconvolution method is able to reproduce almost perfectly the target distribution (MIO close to 1) for large enough dataset sizes, the Fourier-based method always saturates to a suboptimal level even as the number of samples increases, even for low noise levels (large SNR, crosses in Fig. 3). Qualitatively, this is due to the oscillatory features in the distribution produced by the noise during the deconvolution with the FFT method, as a consequence of the oscillatory nature of the Fourier basis functions. In general, it has been proved that in FFT methods the convergence to the actual distribution grows sublinearly with the sample size, with scaling dependencies that make the method unfeasible for practical purposes^16,19,25^. Since Bayesian deconvolution makes use of local basis functions, more parsimonious solutions can be obtained, preventing the degradation of the target distribution due to the noise.

In addition to the direct comparison performed between FFT and Bayesian deconvolutions, we compared the results of the deconvolution against a model that simply fits the convolved data ignoring the convolution problem. A condition to impose on any practical deconvolution algorithm is that the result of the deconvolution is not worse than not doing the decomposition at all. The results of this comparison can be observed in Supplementary Figure S2. The results show that, while the Fourier-based method generally tends to make worse predictions than the null model (fitting without deconvolution), the Bayesian approach consistently generates results virtually as good as when the noise is negligible (SNS=10), and improves systematically the distribution in the high-noise case (SNS=1) over the null model. In conclusion, the Bayesian method that we propose has combined benefits, both qualitatively and quantitatively, over other basis methods as FFT and consistently improves the result (or at least not degrades the quality of the distribution).

### An experimental dataset with external noise

Next we applied our method to an experimental dataset in which the noise distribution is unknown. To generate a ground truth, we use cells with the Brachyury reporter T/Bra::GFP in two media conditions (N2B27 supplemented with 3 *µ*M CHIR99 and DMSO as control). At day 3, for each condition, one of the replicas was treated with 20 nM Green CMFDA dye and incubated for 3 min prior to flow cytometry, and the second one was incubated for 3 min with N2B27 (see Methods). Given that the dye incorporates in the cell cytoplasm and its emission spectrum is similar to the one of GFP, the dye acts as a noise source to the GFP signal coming from the Brachyury reporter. Consequently, we have the following four conditions, with their potential outcomes in terms of the signal measured in the FITC-A channel:

(c1) **Control**: Brachyury expression is minimal, and the signal comes mainly from the intrinsic autofluorescence of the cells.

(c2) **Control**+**CMFDA**: Brachyury expression is minimal, and the signal comes mainly from the CMFDA dye, plus intrinsic autofluorescence of the cells.

(c3) **CHIR99**: Brachyury expression is upregulated, and thus the signal contains both the T/Bra::GFP reporter and the intrinsic autofluorescence of the cells.

(c4) **CHIR99**+**CMFDA**: Brachyury expression is upregulated, and thus the signal contains both the T/Bra::GFP reporter, the signal from the CMFDA dye and the intrinsic autofluorescence of the cells.

The four signal distributions corresponding to these four conditions are shown in Supplementary Fig. S3. We note that the distribution of the signal coming only from the dye cannot be measured independently, given the intrinsic autofluorescence of the cells. Moreover, the dye might be absorbed differently depending on the state of the cells (CHIR99 treatment), and the low concentration of the dye (*∼* nM) might lead to significant cell-to-cell variability on the absorbance of the dye. These features make it a suitable dataset to test the robustness of our method. We first used our method to deconvolve the signal of the dye from the total signal observed in the experimental conditions (c2), using condition (c1) to define the ‘noise’ distribution. The results of the deconvolution are shown in Supplementary Fig. 4, which presents the dye distribution deconvolved from experimental conditions (c1) and (c2). Once the dye distribution was obtained from the deconvolution of conditions (c1) and (c2), we tested the consistency of our approach by considering the dye signal as the noise in condition (c4). Deconvolving the dye distribution from the one measured in (c4) should lead to the distribution obtained in condition (c3), which we can measure experimentally and thus serves as the ground truth in this case. The result can be seen in Fig. 4. Even in this case, where we use a deconvolved dataset for a second deconvolution, we observe that the inferred distribution resulting from our method is in good agreement with the experimentally measured distribution in condition (c3) (MIO = 0.81 *±* 0.04) and improves over the null model that ignores the noise (MIO = 0.46 *±* 0.01).

**FIG. 4.**
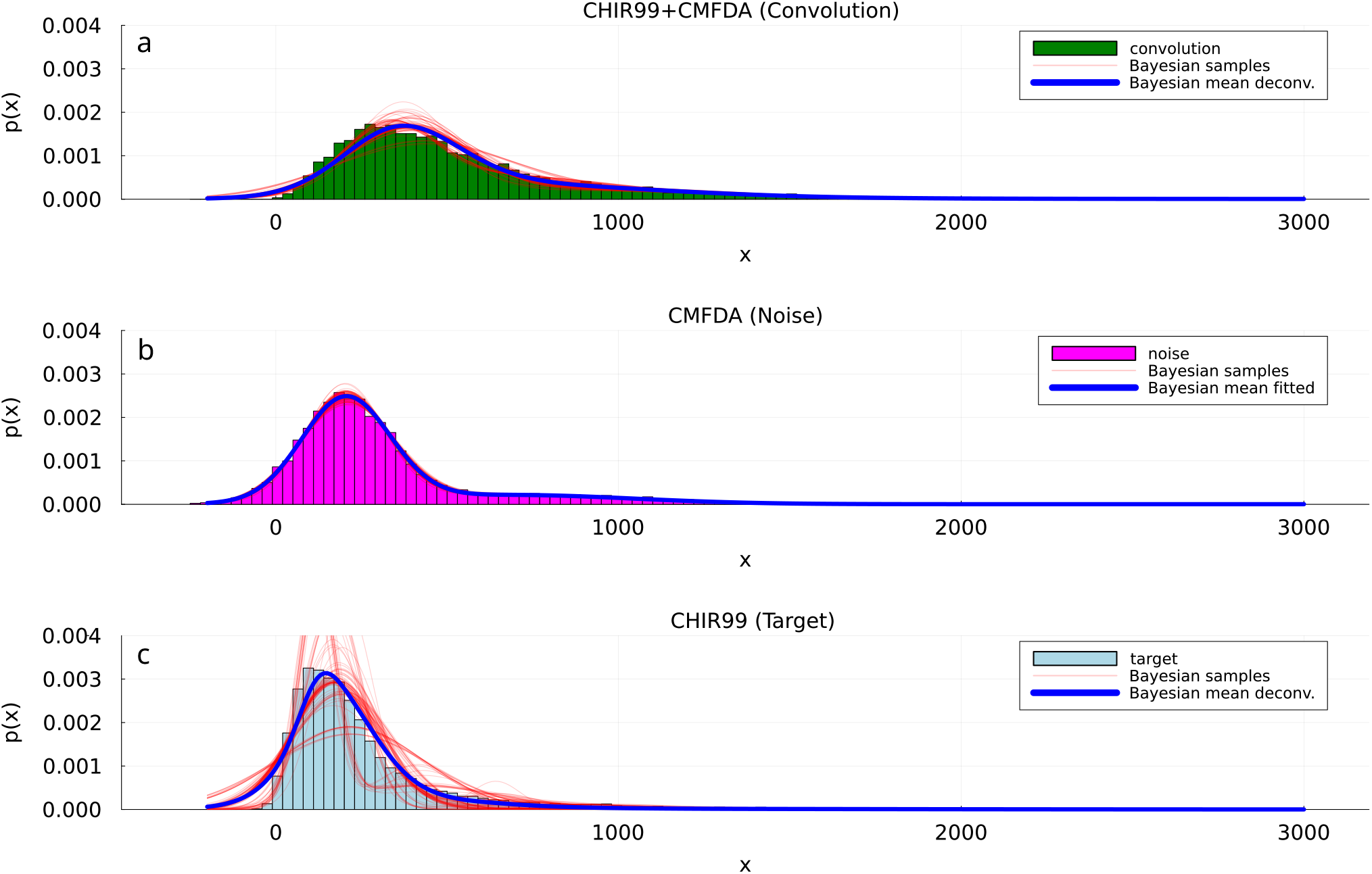
Deconvolution of the CMFDA dye in condition c4 (N2B27+CHIR99+CMFDA). Overlaying the distributions we show realizations of the Bayesian sampling process (red lines) for the three distributions (convolution (a), noise (b) and target (c)) obtained during the fitting. The convolved and target distributions (green and light blue, respectively) come from the real data. The distribution of the dye (in magenta (b)) results from sampling the inferred dye distribution, as described in the text.

Overall, this experiment shows that our method is robust even when our conditions (target and noise independence) are not fully met, and over other fluctuations present in real datasets.

### Multidimensional distributions

Finally, we applied our method in a real flow cytometry dataset with multichannel detection. To that end, we studied the expression of the mesodermal gene Brachyury through a GFP reporter (T/Bra::GFP) in mouse embryonic stem cells (see Methods). In this dataset, two populations are expected to appear: a Brachyury negative population of pluripotent cells, and a positive population capturing the cells exiting from pluripotency towards mesodermal fates. Cells were treated for 24h with 3 *µ*M CHIR99 and 25 ng/mL Activin A to upregulate Brachyury prior to flow cytometry on day 2 (see Methods). The signal of the GFP reporter was our target signal, and it was acquired through the FITC-A channel. In a normal flow cytometry dataset, the signal contains the fluorescence emitted by GFP together with the autofluorescence emitted by the cells at the GFP frequency. The effect of the autofluorescence over the T::GFP signal (target) gives the erroneous impression that Brachyury is expressed in the two populations, when looking directly at the convolved data (light green dots in Figure 5). In other words, it seems there is a non-zero basal expression of the gene in the pluripotent population. When deconvolving the data through the Bayes method, however, the result shows that the pluripotent population peaks actually around zero, as expected since these cells are expected not to show any emission of Brachuyry, which is confirmed by the absence of mRNA in the pluripotent states^31^.

**FIG. 5.**
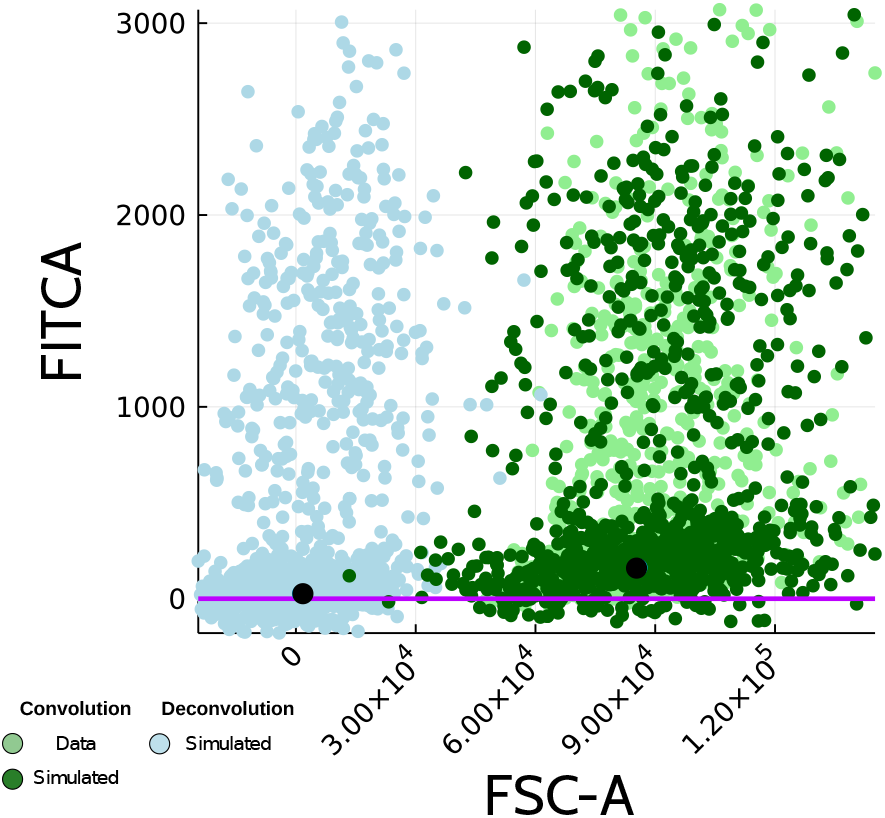
Two-dimensional scatter plot showing a sample of the results of the autofluorescence in a multichannel system. Light and dark green circles are samples of the data from the real data and the fitted convolved distribution, respectively. Light blue circles represent the results of the deconvolution for the two channels. The black dots indicate the mean of the low T:GFP cluster.

Similar problems of non-negative detection have been reported in other highly autofluorescent populations, as in the case of the false positive detection of MHCII in microglia^32^, and the false discovery of Foxp3 expressing cell types that are not T cell lineages^33^.

As can be observed from the other channels (See SSCA in Figure 5 and all the channels in Figure S5), the deconvolution has an effect in removing the expected variance from all the other channels. The left variance may be an effect of the spillover energy that the GFP fluorophore has in the emission spectrum of other channels as the contribution to broad-band channels as FSC and SSC channels (see for example the effect in the FSC-A in Figure 5). The use of deconvolved distributions in single fluorophore samples could improve the estimation of spillover coefficients for sample correction^34^.

In general, the Bayes deconvolution is straightforwardly applicable to real multichannel datasets and its results are consistent with the expected biological observations, showing a clear correction of defects coming from the presence of autofluorescence in the detection channels.

## DISCUSSION

In this paper, we propose a Bayesian approach to obtain flow-cytometry distributions of protein abundance or activity convolved with a known source of noise. The method, which relies on non-parametric Bayesian techniques, is freely available as a Julia package (https://github.com/dsb-lab/scBayesDeconv.jl) and can be used in a straightforward manner in a purely computational way, without the need of dedicated measurement channels or additional laser sources. It only requires measuring the fluorescently labeled and unlabeled cells and, unlike previously proposed deconvolution methods, it provides well-defined and robust probability distributions, described by mixtures of basis functions.

We measure the quality of the results obtained with our method by comparing the deconvolved distributions with known (ground-truth) target distributions using synthetic data and ad hoc experiments with mouse embryonic stem cells. We argue that the use of local basis distributions to describe all the distributions involved in the problem (both measured and unknown), and the corresponding use of a relatively small number of degrees of freedom, leads to an efficient inference. Finally, the Bayesian nature of the method gives rise to a set of candidate target probability distributions. This reflects the natural indeterminacy present in any process corrupted by noise. The ability to express indeterminacy in the solutions is crucial for any real application of a deconvolution algorithm. We further show that the method is robust and consistent even under strong noise in real datasets.

The approach that we propose here is applicable to flow cytometry datasets with multiple channels, which is the current outcome of modern flow cytometers. Furthermore, the method is consistent with already well-established methods for flow cytometry processing, such as the calculation of spillover matrices and the reduction of autofluorescence using additional channels for autofluorescence regression, as discussed in the Supplementary text (Section S1.2). In this sense, our deconvolution approach could be applied to calculate corrected one-fluorophore samples for the calculation of the spillover coefficients, or it could be used after calculating the spillover uncoupling with another method, in order to remove the remaining sources of noise in the dataset.

### LIMITATIONS OF THE STUDY

A potential limitation of the algorithm is the computing time requirements. For *Kn* mixture components of the noise and *Kt* components of the target, a single evaluation of the likelihood function scales as 𝒪 (*Kn ×Kt ×N*), where *N* is the number of measures in the Bayesian model. The current finite mixture implementation is capable of dealing with datasets of 100000 cells with *Kn* = 4 and *Kt* = 6 in about 10 minutes in a desktop with an i7 processor. This efficiency will be sufficient for most part of the analysis, but can be a drawback for the analysis of very large datasets when compared with non-Bayesian methods. The computational time required by the infinite mixture model depends strongly on the parameter *α*, which governs the probability of generating new basis functions, but in general this model is more than an order of magnitude slower than its finite counterpart for the same degree of complexity. Handling large datasets is a well-established problem in Bayesian statistics, and some lines of work have explored solutions to reduce this computational cost, which might be worth exploring in the context of the single-cell Bayesian deconvolution method proposed here^35–37^.

A second potential improvement of the method presented here would be to address the degeneracy under label exchange characteristic of mixture models. Such degeneracy may confound the convergence of the samples to a stationary distribution and the use of the samples for the discovery of clusters, as the same cluster may be assigned to different basis in different samples. A potential solution to this problem would be the implementation of label reassignment methods^38^.

## ACKNOWLEDGEMENTS

We thank the CRG Tissue Engineering Unit and the UPF/CRG Flow Cytometry Unit for continuous support. This work was supported by the Spanish Ministry of Science and Innovation and FEDER, under projects FIS2017-92551-EXP and PID2021-127311NB-I00, by the “Maria de Maeztu” Programme for Units of Excellence in R&D (grant CEX2018-000792-M), and by the Generalitat de Catalunya (ICREA Academia programme). GT is supported by a FPU doctoral fellowship from the Spanish Ministry of Education and Universities (reference FPU18/05091). D.O. acknowledges funding from Juan de la Cierva Incorporación with Project no. IJC2018035298-I, from the Spanish Ministry of Science, Innovation and Universities (MCIU/AEI).

## METHODS

### Synthetic data

For the target distribution we generated samples from three distributions: symmetric bimodal, asymmetric bimodal and skew symmetric distributions (Supplementary Fig. S1). This choice of distributions intends to capture the features present in real datasets, such as the presence of multiple peaks, different cluster sizes, and the generally non-Gaussian character of the data^25^. As for the noise, we generated a set of three different noise datasets containing normal, skewed, and Student’s t distributions (the last two of which exhibit fat tails) (Supplementary Fig. S1), in order to test the flexibility of the method against very dissimilar autofluorescence profiles. In order to check the impact of the noise strength, the convolutions between target and noise were generated at two signal-to-noise ratios (SNR). In our context, we define the SNR as the ratio between target and signal variances for the whole dataset. We chose a case with negligible noise (SNR = 10), and a difficult case where the noise is of the same magnitude as the signal (SNR = 1). We generated sample datasets of different sizes, with 100, 1000 and, 10000 samples. The high range of these values is representative of typical single-cell flow cytometry experiments. The combination of the different target distribution types, noise distribution types, SNRs, and sample sizes generates a collection of 54 datasets with known ground truth.

### Flow cytometry

E14 mouse embryonic stem cells containing a knock-in fluorescence reporter for the mesodermal transcription factor Brachyury, T/Bra::GFP were used^39^. Cells containing the T/Bra::GFP reporter were cultured in ESLif (ESL) medium (KnockOut Dulbecco’s Modified Eagle Medium (DMEM) supplemented with 10% fetal bovine serum (FBS), 1x Non-essential aminoacids (NEEA), 50 U/mL Pen/Strep, 1x GlutaMax, 1x Sodium Pyruvate, 50 *µ*M 2-Mercaptoethanol and leukemia inhibitory factor (LIF)). Cells adhered to 0.1% gelatin-coated (Millipore, ES-006-B) tissue culture-treated 25 cm^2^ (T25 Corning 353108) plates, and were passaged every second day, as previously described^40^. Cells were kept at 37^*°*^C with 5% CO2, and were routinely tested and confirmed to be free from mycoplasma. Flow cytometry experiments were performed as follows: On day 1, cells were trypsinized and seeded into gelatin-coated 6-well plates (Corning, 353224) to a final density of *∼* 10^5^ cells/well in 3 mL ESL media. On day 2, the media was replaced by first washing twice with 3 mL DPBS^+*/*+^ (Phosphate buffered saline containing Mg^++^ and Ca^++^, Sigma, D8662) and then adding NDiff227 media (N2B27) (Takara Bio, #Y40002)^40^ with the appropriate combination of Brachyury activators (3 *µ*M CHIR99, Sigma, SML1046 and/or 25 ng/mL Activin A, Bio-Techne, 338AC-010). The same procedure was followed on day 3. For the control conditions, the corresponding volume of dimethyl sulfoxide (DMSO) was added to the medium. Flow cytometry data was acquired on day 2 or day 3, depending on the type of experiment. For the extrinsic noise experiment, cells were incubated with 1 mL of 20 nM CellTracker Green (CMFDA, Thermofisher) for 3 min prior to trypsinization and flow cytometry. The data was acquired using an LSRIIb flow cytometer, with 2 *×* 10^4^ cells being analyzed per condition. DAPI labelling was used to discard death cells and debris. Furthermore, cell doublets were discarded from the analysis. The readout of protein expression (peak of the signal) was obtained through the FITC-A channel. Measurements were extracted using the FACSDiva software and exported in a Python format for subsequent analysis.

### Software

The software pipeline presented here, including MCMC for finite and infinite normal mixtures, has been implemented as a Julia package (scBayesDeconv.jl). The source code, with manual compilation and installation instructions, as well as full documentation and a notebook with examples of use, can be obtained publicly from GitHub (https://github.com/dsblab/scBayesDeconv.jl).

