## Supplemental file for "Single-cell Bayesian deconvolution"

**S6 Efficiency assessment** **25**
**References** **26**
**Supplementary figures** **27**

### S1 Biological statement of the problem

Flow cytometry provides us with measurements of a target signal $T$ in large numbers of single cells. The target signal is emitted by a fluorophore that reports on the abundance (or activity) of a protein of interest within each cell. This signal is affected by autofluorescence (which can be considered a source of noise) produced by elements of the cell other than the fluorophore. Due to the noise, the total signal $C$ measured by the device is not directly $T$, but

$$C = T + \xi \tag{S1}$$

#### S1.1 Deconvolving signal from noise

We are interested in the case in which we cannot measure both the signal $C$ and the noise $\xi$ independently in the same cell, and thus $T$ in that cell cannot be calculated trivially via Eq. (S1). This limitation is typical of flow cytometry experiments, in which cells can only be measured once. In this case, the only information that can be extracted from the device consists of the distributions of the measured signal, $p_c$, and of the background noise by itself, $p_\xi$ (by measuring cells without fluorophore), over large populations of cells (tens of thousands in a typical flow cytometry run). We can assume the samples to be independent and identically distributed (iid).

If $T$ and $\xi$ and independent of each other, the distributions defined above are related to one another by means of a convolution:

$$p_c(C) = \int_0^\infty \int_0^\infty p_T(T)\, p_\xi(\xi)\, \delta(C - (T + \xi))\, d\xi\, dT = \int_0^\infty p_T(C - \xi)\, p_\xi(\xi)\, d\xi \equiv (p_T * p_\xi)(C) \tag{S2}$$

In what follows, we describe a method to extract the distribution $p_T$ of the target variable $T$ from the observed distributions of $C$ and $\xi$ via a deconvolution of Eq. (S2). The method is applicable to any measurement technique that provides distributions of a signal affected by noise.

#### S1.2 Generalization to multichannel measurements

Flow cytometry systems have multiple detectors to measure light at different emission frequencies. This allows to target different proteins in the same cell with different fluorophores, measuring their emission at either the peak frequency (classical flow cytometry) or in a set of frequencies that define the emission spectrum (spectral flow cytometry). Independently of the method, the emission of a fluorophore extends over the spectrum, and hence it spills over the different channels. Considering the additive effect of the emission of different fluorophore markers on the measured channels, equation (S1) can be more generally stated as

$$c_k = \sum_{j=1}^{N_\mathrm{T}} t_j S_{jk} + \xi_k, \qquad k = 1 \ldots N_\mathrm{ch} \tag{S3}$$

where $c_k$, $t_j$ and $\xi_k$ are realizations of the random variables $C$, $T$ and $\xi$ defined in Eq. (S1) above. The subindex $j$ runs over the $N_{\mathrm{T}}$ fluorophores (one per target), and the subindex $k$ runs over the $N_{\mathrm{ch}}$ channels (so that $\xi_k$ represents the contribution of the autofluorescence to channel $k$). The terms $S_{jk}$ define the spillover matrix $\mathbf{S}$, that quantifies how the fluorophore signals are spread over all the measuring channels.

The spillover matrix can be estimated from single fluorophore controls by using regression methods [1]. To that end, one can perform control experiments in which only one fluorophore is present, and measure the resulting signal in all the channels:

$$c_{ik}^{(j)} = t_{ij}^{(j)} S_{jk} + \xi_{ik} \tag{S4}$$

where the subindex $j$ now corresponds to the only fluorophore present in the system, the superindex $(j)$ indicates that the experiments were performed in the single-fluorophore condition, and $c_{ik}^{(j)}$ denotes a total signal in channel $k$ coming from cell $i$ when only the fluorophore $j$ is present in the system. This set of equations is underdetermined, as we do not know neither the real target signal $t_{ij}^{(j)}$ nor the components $S_{jk}$ of the spillover matrix. However, if we focus for the moment on the channel that corresponds to our single fluorophore ($k = j$) and impose $S_{jj} = 1$, we can write down that $c_{ij}^{(j)} \approx t_{ij}^{(j)}$ (ignoring for now the autofluorescence of that channel). This allows us to establish a set of linear regression problems whose solution enables the estimation of the spillover matrix components for which we have single-fluorophore controls:

$$c_{ik}^{(j)} \approx c_{ij}^{(j)} S_{jk} + \xi_{ik} \tag{S5}$$

The spillover coefficients $S_{jk}$ can then be estimated using robust regression techniques that remove the noise coming from outliers [1].

A similar method can be used to reduce the noise coming from the autofluorescence. To that end, one can define an effective "fluorophore signal" $t_{in}$ coming from the autofluorescence, with an associated additional channel $n$ not linked to a real fluorophore used in the sample. If the signal in this channel is correlated with the emission of the autofluorescence $\xi$ over the other channels, we can decompose the autofluorescence signal at every channel $k$ into a regression term and a noise term:

$$\xi_{ik} \equiv t_{in} S_{nk} + \xi'_{ik} \quad \Longrightarrow \quad c_{ik} = \sum_j t_{ij} S_{jk} + t_{in} S_{nk} + \xi'_{ik} \tag{S6}$$

The coefficients $S_{nk}$ indicate how the autofluorescence signal is spread over the other channels. The fluorophore signals and the autofluorescence "signal" can be grouped in a single term:

$$c_{ik} = \sum_{j=1}^{N_{\mathrm{T}}+1} t_{ij} \bar{S}_{jk} + \xi'_{ik} \tag{S7}$$

where now the sum over $j$ also includes the autofluorescence signal. We can then calculate the spillover coefficients with the method above, by using the additional channel $n$ as an additional "fluorophore control" for the autofluorescence signal. We also note that writing the autofluorescence as a regression problem in Eq. (S6) will lead in general to a reduction of the noise in all channels:

$$\text{Var}(\xi_k') < \text{Var}(\xi_k) \tag{S8}$$

Finally, to obtain the original signal, we can multiply Eq. (S7) by the inverse of the spillover matrix:

$$\mathbf{C}\bar{\mathbf{S}}^{-1} = \mathbf{T} + \boldsymbol{\xi}'\bar{\mathbf{S}}^{-1} \equiv \mathbf{T} + \boldsymbol{\xi}'' \tag{S9}$$

We emphasize that this method requires that the autofluorescence signal at the "noise channel" is sufficiently correlated with its contribution at the other channels, which is not necessarily true *a priori*. Also, Eq. (S9) shows that the need to deconvolve the noise stands even after the application of the spillover and autofluorescence corrections proposed in the literature. In fact, the deconvolution method that we propose in this article is compatible with existing methods of autofluorescence correction such as the regression method reviewed above, which can be performed before applying our method to the corrected system (S9).

#### S2 Mathematical statement of the problem

Since we do not know the exact underlying distributions, we model them as potentially infinite mixtures of normal basis functions. The Gaussian mixture models for the target and the noise distributions can be represented as

$$p(\mathbf{t}|\boldsymbol{\omega}^T, \{\boldsymbol{\mu}^T\}, \{\boldsymbol{\Sigma}^T\}) = \sum_{i=1}^{K_T} \omega_i^T \mathcal{N}\left(\mathbf{t}|\boldsymbol{\mu}_i^T, \boldsymbol{\Sigma}_i^T\right) \tag{S10a}$$

$$p(\boldsymbol{\xi}|\boldsymbol{\omega}^\xi, \{\boldsymbol{\mu}^\xi\}, \{\boldsymbol{\Sigma}^\xi\}) = \sum_{i=1}^{K_\xi} \boldsymbol{\omega}_i^\xi \mathcal{N}\left(\boldsymbol{\xi}|\boldsymbol{\omega}_i^\xi, \boldsymbol{\Sigma}_i^\xi\right), \tag{S10b}$$

where $\mathcal{N}(\boldsymbol{x}|\boldsymbol{\mu}_i, \boldsymbol{\Sigma}_i)$ denotes a normal distribution on the variable vector $\boldsymbol{x}$ (whose components are the different measurement channels), with mean $\boldsymbol{\mu}_i$ and variance $\boldsymbol{\Sigma}_i$. $K_T$ and $K_\xi$ represent the number of bases used to describe each distribution, $\boldsymbol{\omega} = \{\omega_i : i = 1 \ldots K\}$ are their weights, and $\{\boldsymbol{\mu}\} = \{\boldsymbol{\mu}_i : i = 1 \ldots K\}$ and $\{\boldsymbol{\Sigma}\} = \{\boldsymbol{\Sigma}_i : i = 1 \ldots K\}$ are their characteristic parameters. Note that we have removed the subindices from the distributions of the target and the noise, since according to the mixture representation, $p^T$ and $p^\xi$ depend exclusively on the parameters defined above.

The mixture decomposition defined in Eqs. (S10) allows for a flexible and robust representation of unknown and generic distributions. Moreover, exploiting the fact that the convolution of two Gaussian distributions is Gaussian, we have an analytical expression for the distribution of the total variable $\mathbf{C}$:

$$p(\boldsymbol{c}|\boldsymbol{\phi}^T, \boldsymbol{\phi}^\xi) = \sum_{i=1}^{K_T} \sum_{j=1}^{K_\xi} \omega_j^T \omega_j^\xi \mathcal{N}\left(\boldsymbol{c} \,\middle|\, \boldsymbol{\mu}_i^T + \boldsymbol{\mu}_j^\xi, \boldsymbol{\Sigma}_i^T + \boldsymbol{\Sigma}_j^\xi\right) \tag{S11}$$

where $\boldsymbol{\phi}^T = \{\boldsymbol{\omega}^T, \boldsymbol{\mu}^T, \boldsymbol{\Sigma}^T\}$ and $\boldsymbol{\phi}^\xi = \{\boldsymbol{\omega}^\xi, \boldsymbol{\mu}^\xi, \boldsymbol{\Sigma}^\xi\}$ represent all the parameters of the target and noise distributions, from which we can parametrize the distribution of the total measured signal. We note that the combination of normal basis function $\mathcal{N}$ in Eq. (S11) above is invariant under changes in the individual basis functions of $T$ and $\xi$, provided the sums of means, $\boldsymbol{\mu}^T + \boldsymbol{\mu}^\xi$, and variances, $\boldsymbol{\Sigma}^T + \boldsymbol{\Sigma}^\xi$, are constant.

As discussed in the main text, according to Bayes' rule, the posterior distribution that represents the probability of the parameters given the data is

$$p(\boldsymbol{\phi}^T, \boldsymbol{\phi}^\xi | \{\mathbf{c}\}, \{\boldsymbol{\xi}\}) \propto \Big[p(\{\mathbf{c}\}|\boldsymbol{\phi}^T, \boldsymbol{\phi}^\xi) p(\{\boldsymbol{\xi}\}|\boldsymbol{\phi}^\xi)\Big] \Big[p(\boldsymbol{\phi}^\xi) p(\boldsymbol{\phi}^T)\Big], \tag{S12}$$

where $\{\mathbf{c}\} = \{\boldsymbol{c}_i : i = 1 \ldots N_c\}$ and $\{\boldsymbol{\xi}\} = \{\boldsymbol{\xi}_i : i = 1 \ldots N_\xi\}$ represent the samples of the total signal and noise, respectively, with the different components of $\boldsymbol{c}_i$ and $\boldsymbol{\xi}_i$ correspond to the different channels of the cytometer. In Eq. (S12), the first bracket on the right-hand side corresponds to the likelihood, namely the joint probability of observing the data given the parameters, which we can redefine as

$$\mathcal{L} = p(\{\mathbf{c}\}|\boldsymbol{\phi}^T, \boldsymbol{\phi}^\xi) p(\{\boldsymbol{\xi}\}|\boldsymbol{\phi}^\xi) = \prod_i^{N_c} p(\boldsymbol{c}_i|\boldsymbol{\phi}^T, \boldsymbol{\phi}^\xi) \prod_j^{N_\xi} p(\boldsymbol{\xi}_j|\boldsymbol{\phi}^\xi), \tag{S13}$$

The form of the likelihood (S13) breaks the symmetry between the target and noise signals, and thus lifts the above-mentioned degeneracy between their parameters exhibited by Eq. (S11). The second bracket in the right-hand side of Eq. (S12), in turn, corresponds to the prior distributions of all the parameters of the problem, which we define in what follows.

##### S2.1 Approximated decomposition of the posterior into two separate problems

It is worth noting that the posterior distribution (S12) can be decomposed as follows:

$$p(\boldsymbol{\phi}^T, \boldsymbol{\phi}^\xi | \{\boldsymbol{c}\}, \{\boldsymbol{\xi}\}) = p(\boldsymbol{\phi}^T | \boldsymbol{\phi}^\xi, \{\boldsymbol{c}\}) p(\boldsymbol{\phi}^\xi | \{\boldsymbol{c}\}, \{\boldsymbol{\xi}\}) \tag{S14}$$

where we have removed the dependency of the noise signal in the first term of the right-hand side, since only the total signal $\boldsymbol{c}$ defines the parameters of the target distribution, as can be seen from the likelihood

(S13). The second term, on the other hand, is conditioned by both the noise and the total signal. Since we usually have as much data for the noise signal $\{\boldsymbol{\xi}\}$ as for the signal of interest $\{\boldsymbol{c}\}$, we can approximately consider that the posterior distribution of the noise mixture parameter $\boldsymbol{\phi}^{\xi}$ is well represented by the noise data alone:

$$p(\boldsymbol{\phi}^T, \boldsymbol{\phi}^{\xi}|\{\mathbf{c}\}, \{\boldsymbol{\xi}\}) \approx p(\boldsymbol{\phi}^T|\boldsymbol{\phi}^{\xi}, \{\mathbf{c}\})p(\boldsymbol{\phi}^{\xi}|\{\boldsymbol{\xi}\}) \tag{S15}$$

With this approximation, the problem can be decomposed in two separate subproblems: first, finding the probability distribution of the parameters $\boldsymbol{\phi}^{\xi}$ of the noise mixture; and second, finding the distribution of the parameters $\boldsymbol{\phi}^T$ of the convolution mixture, conditioned on the noise mixture parameters.

In the following section, we go over the mathematical details of the probability distributions that will allow us to sample from the posterior distribution.

### S3 Relevant probability distributions

In this section, we derive the main expressions that we need to sample our model. Our aim is to have a self-contained derivation of the sampling process of the posterior of a Gaussian normal mixture.

#### S3.1 Multivariate normal distribution with unknown mean and error

Here we derive the main results of interest on multivariate normal distributions, which we will use when sampling the posterior distribution using normal bases.

##### S3.1.1 Likelihood

The multivariate normal distribution for a set of independent identical samples (iid) $\{\boldsymbol{x}\}$ has the form, up to a scaling parameter,

$$\mathcal{N}(\{\boldsymbol{x}\}; \boldsymbol{\mu}, \boldsymbol{\Sigma}) \propto \det[\boldsymbol{\tau}]^{n/2} \exp\left\{-\frac{1}{2}\left[\sum_{i=1}^{n}(\boldsymbol{x}_i - \boldsymbol{\mu})^T\boldsymbol{\tau}(\boldsymbol{x}_i - \boldsymbol{\mu})\right]\right\} \tag{S16}$$

where $\mu$ is the mean and $\boldsymbol{\tau}$ is the precision parameter and $n$ is the number of cells being measured. The precision parameter relates to the covariance matrix as

$$\boldsymbol{\tau} = \boldsymbol{\Sigma}^{-1} \tag{S17}$$

It is convenient to rearrange the distribution to make explicit the dependency of the summary statistics:

$$\begin{aligned}\mathcal{N}(\{\boldsymbol{x}\};\boldsymbol{\mu},\boldsymbol{\Sigma}) &\propto \det[\boldsymbol{\tau}]^{n/2}\exp\left\{-\frac{1}{2}\left[\sum_i(\boldsymbol{x}_i-\boldsymbol{\mu})^T\boldsymbol{\tau}(\boldsymbol{x}_i-\boldsymbol{\mu})\right]\right\}\\ &\propto \det[\boldsymbol{\tau}]^{n/2}\exp\left\{-\frac{1}{2}\left[\sum_i(\boldsymbol{x}_i-\bar{\boldsymbol{x}})^T\boldsymbol{\tau}(\boldsymbol{x}_i-\bar{\boldsymbol{x}})+n(\bar{\boldsymbol{x}}-\boldsymbol{\mu})^T\boldsymbol{\tau}(\bar{\boldsymbol{x}}-\boldsymbol{\mu})\right]\right\}\\ &\propto \det[\boldsymbol{\tau}]^{n/2}\exp\left\{-\frac{1}{2}\left[n\mathrm{Tr}\left[\boldsymbol{S}_x^2\boldsymbol{\tau}\right]+n(\bar{\boldsymbol{x}}-\boldsymbol{\mu})^T\boldsymbol{\tau}(\bar{\boldsymbol{x}}-\boldsymbol{\mu})\right]\right\} \end{aligned}\tag{S18}$$

where the summary statistics are the mean,

$$\bar{\boldsymbol{x}}=\frac{1}{n}\sum_i\boldsymbol{x}_i \tag{S19}$$

and the covariance,

$$\boldsymbol{S}_x^2=\frac{1}{n}\sum_i(\boldsymbol{x}_i-\bar{\boldsymbol{x}})(\boldsymbol{x}_i-\bar{\boldsymbol{x}})^T \tag{S20}$$

###### S3.1.2 Conjugate prior

A convenient prior for the multivariate distribution is a conjugate prior that allows us to obtain the analytic form of the other distributions:

$$p(\boldsymbol{\mu}|\boldsymbol{\Sigma})=\mathcal{N}(\boldsymbol{\mu};\boldsymbol{\mu}_0,\kappa_0\boldsymbol{\Sigma}) \tag{S21}$$

$$p(\boldsymbol{\Sigma})=\mathcal{W}^{-1}(\boldsymbol{\Sigma};\boldsymbol{\Sigma}_0,\nu_0) \tag{S22}$$

where $\mathcal{W}^{-1}$ is the inverse Wishart distribution. The hyperparameter $\kappa_0$ represents our confidence in the estimation of the mean: the higher $\kappa_0$ is, the closer to the mean we will be.

The inverse Wishart distribution has the following shape, up to scaling terms:

$$\mathcal{W}^{-1}(\boldsymbol{X};\boldsymbol{\Sigma},\nu)\propto\frac{\det[\boldsymbol{\Sigma}]^{(\nu)/2}}{\det[\boldsymbol{X}]^{(\nu+p+1)/2}}e^{-\mathrm{Tr}[\boldsymbol{\Sigma}\boldsymbol{X}^{-1}]/2} \tag{S23}$$

where $p$ is the number of dimensions. The parameter $\boldsymbol{\Sigma}$ is the correlation matrix and $\nu$ represents our confidence in the estimation of the $\boldsymbol{\Sigma}$ correlation matrix.

##### S3.1.3 Posterior distribution

The posterior distribution will have the following shape:

$$
\begin{aligned}
p(\boldsymbol{\mu}, \boldsymbol{\Sigma}|\{\boldsymbol{x}\}) \propto\ & p(\{\boldsymbol{x}\}|\boldsymbol{\mu}, \boldsymbol{\Sigma}) p(\boldsymbol{\mu}|\boldsymbol{\Sigma}, \kappa_0, \boldsymbol{\mu}_0) p(\boldsymbol{\Sigma}|\boldsymbol{\Sigma}_0, \nu_0) \\
& \propto \det[\boldsymbol{\tau}]^{n/2} \exp\left\{-\frac{1}{2}\left[n\mathrm{Tr}\left[\boldsymbol{S}_x^2\boldsymbol{\tau}\right] + n(\bar{\boldsymbol{x}} - \boldsymbol{\mu})^T\boldsymbol{\tau}(\bar{\boldsymbol{x}} - \boldsymbol{\mu})\right]\right\} \times \\
\det[\boldsymbol{\tau}]^{1/2} \exp & \left\{-\frac{1}{2}\left[\kappa_0(\boldsymbol{\mu} - \boldsymbol{\mu}_0)^T\boldsymbol{\tau}(\boldsymbol{\mu} - \boldsymbol{\mu}_0)\right]\right\} \det[\boldsymbol{\tau}]^{(\nu_0+p+1)/2} \exp\left\{-\frac{1}{2}\left[\mathrm{Tr}\left[\boldsymbol{\Sigma}_0\boldsymbol{\tau}\right]\right]\right\} \qquad \text{(S24)}
\end{aligned}
$$

We can use the posterior distribution to obtain different distributions of relevance.

##### S3.1.4 Conditional distribution: mean

Retaining the terms involving the mean $\boldsymbol{\mu}$, the conditional distribution of the mean takes the form of a multivariate distribution:

$$
\begin{aligned}
p(\boldsymbol{\mu}|\boldsymbol{\Sigma}, \{\boldsymbol{x}\}) &\propto \exp\left\{-\frac{1}{2}\left[n(\bar{\boldsymbol{x}} - \boldsymbol{\mu})^T\boldsymbol{\tau}(\bar{\boldsymbol{x}} - \boldsymbol{\mu}) + \kappa_0(\boldsymbol{\mu} - \boldsymbol{\mu}_0)^T\boldsymbol{\tau}(\boldsymbol{\mu} - \boldsymbol{\mu}_0)\right]\right\} \\
&\propto \exp\left\{-\frac{1}{2}\left[-2(n\bar{\boldsymbol{x}}^T + \kappa_0\boldsymbol{\mu}_0)\boldsymbol{\tau}\boldsymbol{\mu} + (n + \kappa_0)\boldsymbol{\mu}^T\boldsymbol{\tau}\boldsymbol{\mu}\right]\right\} \\
&\propto \exp\left\{-\frac{1}{2}\left[(\boldsymbol{\mu} - \tilde{\boldsymbol{\mu}})^T\tilde{\boldsymbol{\tau}}(\boldsymbol{\mu} - \tilde{\boldsymbol{\mu}})\right]\right\} \quad \propto \mathcal{N}(\boldsymbol{\mu}|\tilde{\boldsymbol{\mu}}, \tilde{\boldsymbol{\tau}}) \qquad \text{(S25)}
\end{aligned}
$$

where in the last step we completed the squares. This multivariate normal distribution has effective parameters

$$
\tilde{\boldsymbol{\mu}} = \frac{n\bar{\boldsymbol{x}} + \kappa_0\boldsymbol{\mu}_0}{n + \kappa_0}, \qquad \tilde{\boldsymbol{\tau}} = (n + \kappa_0)\boldsymbol{\tau} \qquad \text{(S26)}
$$

where $\tilde{\boldsymbol{\mu}}$ is a weighted version between the mean statistic and the prior mean, and $\tilde{\boldsymbol{\tau}}$ is an effective precision parameter. The $\kappa_0$ factor indicates how close we are from the prior mean.

##### S3.1.5 Conditional distribution: covariance

Retaining the terms involving the covariance matrix (and its inverse, the precision matrix), the conditional distribution can be shown to take the form of an inverse Wishart distribution:

$$
\begin{aligned}
&p(\mathbf{\Sigma}|\boldsymbol{\mu},\{\boldsymbol{x}\}) \propto \\
&\det[\boldsymbol{\tau}]^{(n+\nu_0+p+2)/2}\exp\left\{-\frac{1}{2}\left[n\mathrm{Tr}\left[\boldsymbol{S}_x^2\boldsymbol{\tau}\right]+n(\bar{\boldsymbol{x}}-\boldsymbol{\mu})^T\boldsymbol{\tau}(\bar{\boldsymbol{x}}-\boldsymbol{\mu})+\kappa_0(\boldsymbol{\mu}-\boldsymbol{\mu}_0)^T\boldsymbol{\tau}(\boldsymbol{\mu}-\boldsymbol{\mu}_0)+\mathrm{Tr}\left[\mathbf{\Sigma}_0\boldsymbol{\tau}\right]\right]\right\} \\
&\propto \det[\boldsymbol{\tau}]^{(n+\nu_0+p+2)/2}\exp\left\{-\frac{1}{2}\left[\mathrm{Tr}\left[\left(n\boldsymbol{S}_x^2\boldsymbol{\tau}+n(\bar{\boldsymbol{x}}-\boldsymbol{\mu})(\bar{\boldsymbol{x}}-\boldsymbol{\mu})^T+\kappa_0(\boldsymbol{\mu}-\boldsymbol{\mu}_0)(\boldsymbol{\mu}-\boldsymbol{\mu}_0)^T+\mathbf{\Sigma}_0\right)\boldsymbol{\tau}\right]\right]\right\} \\
&\propto \det[\boldsymbol{\tau}]^{(\tilde{n}+p+1)/2}\exp\left\{-\frac{1}{2}\left[\mathrm{Tr}\left[\tilde{\mathbf{\Sigma}}\boldsymbol{\tau}\right]\right]\right\} \quad \propto \mathcal{W}^{-1}(\mathbf{\Sigma}|\tilde{\mathbf{\Sigma}},\tilde{n})
\end{aligned} \tag{S27}
$$

where the effective parameters are

$$\tilde{n} = n + \nu_0 + 1 \tag{S28}$$

$$\tilde{\mathbf{\Sigma}} = n\boldsymbol{S}_x^2 + n(\bar{\boldsymbol{x}}-\boldsymbol{\mu})(\bar{\boldsymbol{x}}-\boldsymbol{\mu})^T + \kappa_0(\boldsymbol{\mu}-\boldsymbol{\mu}_0)(\boldsymbol{\mu}-\boldsymbol{\mu}_0)^T + \mathbf{\Sigma}_0 \tag{S29}$$

In this last expression, it is worth noting that each term represents a different kind of uncertainty: the first term is the uncertainty coming from the mean statistic, the second corresponds to the uncertainty with which the mean statistic represents the actual mean, the third term is the uncertainty coming from the prior mean, and the last term is the uncertainty coming from the prior itself.

##### S3.1.6 Marginal distribution: covariance

One additional distribution that we will need is the marginal distribution of the variance:

$$
\begin{aligned}
p(\mathbf{\Sigma}|\{\boldsymbol{x}\}) &= \int p(\mathbf{\Sigma},\boldsymbol{\mu}|\{\boldsymbol{x}_i\}_i)d\boldsymbol{\mu} \propto \int \det[\boldsymbol{\tau}]^{n/2}\exp\left\{-\frac{1}{2}\left[n\mathrm{Tr}\left[\boldsymbol{S}_x^2\boldsymbol{\tau}\right]+n(\bar{\boldsymbol{x}}-\boldsymbol{\mu})^T\boldsymbol{\tau}(\bar{\boldsymbol{x}}-\boldsymbol{\mu})\right]\right\} \\
&\times \det[\boldsymbol{\tau}]^{1/2}\exp\left\{-\frac{1}{2}\left[\kappa_0(\boldsymbol{\mu}-\boldsymbol{\mu}_0)^T\boldsymbol{\tau}(\boldsymbol{\mu}-\boldsymbol{\mu}_0)\right]\right\}\det[\boldsymbol{\tau}]^{(\nu_0+p+1)/2}\exp\left\{-\frac{1}{2}\left[\mathrm{Tr}\left[\mathbf{\Sigma}_0\boldsymbol{\tau}\right]\right]\right\}d\boldsymbol{\mu} \\
&\propto \det[\boldsymbol{\tau}]^{(n+\nu_0+p+2)/2}\exp\left\{-\frac{1}{2}\left[n\mathrm{Tr}\left[\boldsymbol{S}_x^2\boldsymbol{\tau}\right]+\mathrm{Tr}\left[\mathbf{\Sigma}_0\boldsymbol{\tau}\right]\right]\right\} \\
&\times \left[\int \exp\left\{-\frac{1}{2}\left[\underbrace{\kappa_0(\boldsymbol{\mu}-\boldsymbol{\mu}_0)^T\boldsymbol{\tau}(\boldsymbol{\mu}-\boldsymbol{\mu}_0)+n(\bar{\boldsymbol{x}}-\boldsymbol{\mu})^T\boldsymbol{\tau}(\bar{\boldsymbol{x}}-\boldsymbol{\mu})}_{(*)}d\boldsymbol{\mu}\right]\right\}\right]
\end{aligned} \tag{S30}
$$

Reorganizing the elements in $(*)$,

$$(*) = (\kappa_0 + n)\boldsymbol{\mu}^T\boldsymbol{\tau}\boldsymbol{\mu} - 2\boldsymbol{\mu}^T\boldsymbol{\tau}(\kappa_0\boldsymbol{\mu}_0 + n\bar{\boldsymbol{x}}) + (n\bar{\boldsymbol{x}}^T\boldsymbol{\tau}\bar{\boldsymbol{x}} + \kappa_0\boldsymbol{\mu}_0\boldsymbol{\tau}\boldsymbol{\mu}_0) \tag{S31}$$

$$= (\boldsymbol{\mu} - \tilde{\boldsymbol{\mu}})^T\tilde{\boldsymbol{\tau}}(\boldsymbol{\mu} - \tilde{\boldsymbol{\mu}}) \underbrace{-\tilde{\boldsymbol{\mu}}^T\tilde{\boldsymbol{\tau}}\tilde{\boldsymbol{\mu}} + (n\bar{\boldsymbol{x}}^T\boldsymbol{\tau}\bar{\boldsymbol{x}} + \kappa_0\boldsymbol{\mu}_0\boldsymbol{\tau}\boldsymbol{\mu}_0),}_{(**)} \tag{S32}$$

where we have defined,

$$\tilde{\boldsymbol{\mu}} = \frac{\kappa_0\boldsymbol{\mu}_0 + n\bar{\boldsymbol{x}}}{\kappa_0 + n} \tag{S33}$$

$$\tilde{\boldsymbol{\tau}} = (\kappa_0 + n)\boldsymbol{\tau}, \tag{S34}$$

we can further regroup the part in $(**)$.

$$(**) = \frac{-\kappa_0^2\boldsymbol{\mu}_0^T\boldsymbol{\tau}\boldsymbol{\mu}_0 - n^2\bar{\boldsymbol{x}}^T\boldsymbol{\tau}\bar{\boldsymbol{x}} - 2\kappa_0 n\boldsymbol{\mu}_0^T\boldsymbol{\tau}\bar{\boldsymbol{x}} + n^2\bar{\boldsymbol{x}}^T\boldsymbol{\tau}\bar{\boldsymbol{x}} + \kappa_0^2\boldsymbol{\mu}_0\boldsymbol{\tau}\boldsymbol{\mu}_0 + n\kappa_0\bar{\boldsymbol{x}}^T\boldsymbol{\tau}\bar{\boldsymbol{x}} + \kappa_0 n\boldsymbol{\mu}_0\boldsymbol{\tau}\boldsymbol{\mu}_0}{\kappa_0 + n}$$

$$\frac{\kappa_0 n(\bar{\boldsymbol{x}}^T\boldsymbol{\tau}\bar{\boldsymbol{x}} - 2\boldsymbol{\mu}_0^T\boldsymbol{\tau}\bar{\boldsymbol{x}} + \boldsymbol{\mu}_0\boldsymbol{\tau}\boldsymbol{\mu}_0)}{\kappa_0 + n} = \frac{\kappa_0 n}{\kappa_0 + n}(\bar{\boldsymbol{x}} - \boldsymbol{\mu}_0)^T\boldsymbol{\tau}(\bar{\boldsymbol{x}} - \boldsymbol{\mu}_0) \tag{S35}$$

Inserting these terms

$$p(\boldsymbol{\Sigma}|\{\boldsymbol{x}\}) \propto \det[\boldsymbol{\tau}]^{(n+\nu_0+p+2)/2} \exp\left\{-\frac{1}{2}\left[\mathrm{Tr}\left[\left(n\boldsymbol{S}_x^2 + \boldsymbol{\Sigma}_0 + \frac{\kappa_0 n}{\kappa_0 + n}(\bar{\boldsymbol{x}} - \boldsymbol{\mu}_0)(\bar{\boldsymbol{x}} - \boldsymbol{\mu}_0)^T\right)\boldsymbol{\tau}\right]\right]\right\}$$
$$\times \left[\int \exp\left\{-\frac{1}{2}\left[(\boldsymbol{\mu} - \tilde{\boldsymbol{\mu}})^T\tilde{\boldsymbol{\tau}}(\boldsymbol{\mu} - \tilde{\boldsymbol{\mu}})\right]\right\} d\boldsymbol{\mu}\right] \tag{S36}$$

now we can calculate the integral as a multivariate normal:

$$p(\boldsymbol{\Sigma}|\{\boldsymbol{x}\}) \propto$$
$$\propto \det[\boldsymbol{\tau}]^{(n+\nu_0+p+2)/2} \exp\left\{-\frac{1}{2}\left[\mathrm{Tr}\left[\left(n\boldsymbol{S}_x^2 + \boldsymbol{\Sigma}_0 + \frac{\kappa_0 n}{\kappa_0 + n}(\bar{\boldsymbol{x}} - \boldsymbol{\mu}_0)(\bar{\boldsymbol{x}} - \boldsymbol{\mu}_0)^T\right)\boldsymbol{\tau}\right]\right]\right\} \det[\tilde{\boldsymbol{\tau}}]^{-1/2}$$
$$\propto \det[\boldsymbol{\tau}]^{(n+\nu_0+p+1)/2} \exp\left\{-\frac{1}{2}\left[\mathrm{Tr}\left[\tilde{\boldsymbol{\Sigma}}\boldsymbol{\tau}\right]\right]\right\} \propto \mathcal{W}^{-1}(\boldsymbol{\Sigma}|\tilde{\boldsymbol{\Sigma}}, \tilde{n}) \tag{S37}$$

where in the last step we use the fact that $\det\tilde{\boldsymbol{\tau}} \propto \det\boldsymbol{\tau}$ and the effective parameters of the inverse Wishart distribution are

$$\tilde{n} = n + \nu_0 + 1 \tag{S38}$$

$$\tilde{\boldsymbol{\Sigma}} = n\boldsymbol{S}_x^2 + \frac{\kappa_0 n}{\kappa_0 + n}(\bar{\boldsymbol{x}} - \boldsymbol{\mu}_0)(\bar{\boldsymbol{x}} - \boldsymbol{\mu}_0)^T + \boldsymbol{\Sigma}_0 \tag{S39}$$

###### S3.1.7 Posterior predictive distribution

One last distribution of interest is the probability of a new data point given the already observed set of observations:

$$
\begin{aligned}
p(\boldsymbol{y}|\{\boldsymbol{x}\}) = \iint p(\boldsymbol{y}|\boldsymbol{\mu},\boldsymbol{\Sigma})p(\boldsymbol{\mu},\boldsymbol{\Sigma}|\{\boldsymbol{x}_i\}_i)d\boldsymbol{\mu}d\boldsymbol{\Sigma} \quad &\propto \iint \det[\boldsymbol{\tau}]^{(n+\nu_0+p+3)/2} \\
&\times \exp\left\{-\frac{1}{2}\left[\underbrace{(\boldsymbol{y}-\boldsymbol{\mu})^T\boldsymbol{\tau}(\boldsymbol{y}-\boldsymbol{\mu}) + n(\bar{\boldsymbol{x}}-\boldsymbol{\mu})^T\boldsymbol{\tau}(\bar{\boldsymbol{x}}-\boldsymbol{\mu}) + \kappa_0(\boldsymbol{\mu}-\boldsymbol{\mu}_0)^T\boldsymbol{\tau}(\boldsymbol{\mu}-\boldsymbol{\mu}_0)}_{(*)}\right]\right\} \\
&\times \exp\left\{-\frac{1}{2}\left[\mathrm{Tr}\left[(n\boldsymbol{S}_x^2+\boldsymbol{\Sigma}_0)\boldsymbol{\tau}\right]\right]\right\} d\boldsymbol{\mu}d\boldsymbol{\Sigma} \qquad \text{(S40)}
\end{aligned}
$$

We can rearrange the elements in $(*)$, completing squares as we did when calculating the mean conditional distribution (sectionS3.1.4):

$$
(*) = \underbrace{n\bar{\boldsymbol{x}}^T\boldsymbol{\tau}\bar{\boldsymbol{x}} + \boldsymbol{y}^T\boldsymbol{\tau}\boldsymbol{y} + \kappa_0\boldsymbol{\mu}_0^T\boldsymbol{\tau}\boldsymbol{\mu}_0 - \tilde{\boldsymbol{\mu}}\tilde{\boldsymbol{\tau}}\tilde{\boldsymbol{\mu}}^{\mu}}_{(**)} + (\boldsymbol{\mu}-\tilde{\boldsymbol{\mu}}^{\mu})^T\tilde{\boldsymbol{\tau}}(\boldsymbol{\mu}-\tilde{\boldsymbol{\mu}}^{\mu}), \qquad \text{(S41)}
$$

where we have retained all the terms involving $\boldsymbol{\tau}$ and $\boldsymbol{\mu}$ this time, as they are necessary for integrating out. The effective parameters are now

$$
\tilde{\boldsymbol{\mu}}^{\mu} = \frac{n\bar{\boldsymbol{x}} + \kappa_0\boldsymbol{\mu}_0 + \boldsymbol{y}}{n+\kappa_0+1} \qquad \text{(S42)}
$$

$$
\tilde{\boldsymbol{\tau}}^{\mu} = (n+\kappa_0+1)\boldsymbol{\tau} \qquad \text{(S43)}
$$

We can further rearrange the terms in $(**)$ to obtain an expression with the $y$ in quadrature:

$$
\begin{aligned}
(**) &= n\bar{\boldsymbol{x}}^T\boldsymbol{\tau}\bar{\boldsymbol{x}} + \boldsymbol{y}^T\boldsymbol{\tau}\boldsymbol{y} + \kappa_0\boldsymbol{\mu}_0^T\boldsymbol{\tau}\boldsymbol{\mu}_0 - \frac{1}{n+\kappa_0+1}(n\bar{\boldsymbol{x}}+\kappa_0\boldsymbol{\mu}_0+\boldsymbol{y})^T\boldsymbol{\tau}(n\bar{\boldsymbol{x}}+\kappa_0\boldsymbol{\mu}_0+\boldsymbol{y}) \\
&= \frac{1}{n+\kappa_0+1}\left[(n\bar{\boldsymbol{x}}^T\boldsymbol{\tau}\bar{\boldsymbol{x}} + \boldsymbol{y}^T\boldsymbol{\tau}\boldsymbol{y} + \kappa_0\boldsymbol{\mu}_0^T\boldsymbol{\tau}\boldsymbol{\mu}_0)(n+\kappa_0+1) - (n\bar{\boldsymbol{x}}+\kappa_0\boldsymbol{\mu}_0+\boldsymbol{y})^T\boldsymbol{\tau}(n\bar{\boldsymbol{x}}+\kappa_0\boldsymbol{\mu}_0+\boldsymbol{y})\right] \\
&= \frac{1}{n+\kappa_0+1}\left[n(\boldsymbol{y}-\bar{\boldsymbol{x}})^T\boldsymbol{\tau}(\boldsymbol{y}-\bar{\boldsymbol{x}}) + \kappa_0(\boldsymbol{y}-\boldsymbol{\mu}_0)^T\boldsymbol{\tau}(\boldsymbol{y}-\boldsymbol{\mu}_0) + n\kappa_0(\boldsymbol{\mu}_0-\bar{\boldsymbol{x}})^T\boldsymbol{\tau}(\boldsymbol{\mu}_0-\bar{\boldsymbol{x}})\right] \\
&= \frac{(n+\kappa_0)}{n+\kappa_0+1}(\boldsymbol{y}-\tilde{\boldsymbol{\mu}}^y)^T\boldsymbol{\tau}(\boldsymbol{y}-\tilde{\boldsymbol{\mu}}^y) + \frac{(\frac{n\kappa_0}{n+\kappa_0}+n\kappa_0)}{n+\kappa_0+1}(\boldsymbol{\mu}_0-\bar{\boldsymbol{x}})^T\boldsymbol{\tau}(\boldsymbol{\mu}_0-\bar{\boldsymbol{x}}) \\
&= m^0(\boldsymbol{y}-\tilde{\boldsymbol{\mu}}^y)^T\boldsymbol{\tau}(\boldsymbol{y}-\tilde{\boldsymbol{\mu}}^y) + m^1(\boldsymbol{\mu}_0-\bar{\boldsymbol{x}})^T\boldsymbol{\tau}(\boldsymbol{\mu}_0-\bar{\boldsymbol{x}}) \qquad \text{(S44)}
\end{aligned}
$$

where in the second to the third step we group by quadrature and complete squares to group the terms with $y$, and then group by quadrature the terms on the left. The effective parameters are

$$
\tilde{\boldsymbol{\mu}}^y = \frac{(n\bar{\boldsymbol{x}}+\kappa_0\boldsymbol{\mu}_0)}{n+\kappa_0}, \qquad m^0 = \frac{n+\kappa_0}{n+\kappa_0+1}, \qquad m^1 = \frac{\frac{n\kappa_0}{n+\kappa_0}+n\kappa_0}{n+\kappa_0+1} \qquad \text{(S45)}
$$

We can now insert the obtained term in the original expression:

$$
\begin{aligned}
p(\boldsymbol{y}|\{\boldsymbol{x}\}) \propto & \iint \det[\boldsymbol{\tau}]^{(n+\nu_0+p+3)/2} \times \\
& \exp\left\{-\frac{1}{2}\left[m^0(\boldsymbol{y}-\tilde{\boldsymbol{\mu}}^y)^T\boldsymbol{\tau}(\boldsymbol{y}-\tilde{\boldsymbol{\mu}}^y)+m^1(\boldsymbol{\mu}_0-\bar{\boldsymbol{x}})^T\boldsymbol{\tau}(\boldsymbol{\mu}_0-\bar{\boldsymbol{x}})+\mathrm{Tr}\left[n\boldsymbol{S}_x^2\boldsymbol{\tau}\right]+\mathrm{Tr}\left[\boldsymbol{\Sigma}_0\boldsymbol{\tau}\right]\right]\right\}\times \\
& \exp\left\{-\frac{1}{2}\left[(\boldsymbol{\mu}-\tilde{\boldsymbol{\mu}}^\mu)^T\tilde{\boldsymbol{\tau}}^\mu(\boldsymbol{\mu}-\tilde{\boldsymbol{\mu}}^\mu)\right]\right\}d\boldsymbol{\mu}d\boldsymbol{\Sigma} \\
& = \int \det[\boldsymbol{\tau}]^{(n+\nu_0+p+3)/2}\exp\left\{-\frac{1}{2}\left[\mathrm{Tr}\left[\tilde{\boldsymbol{\Sigma}}\boldsymbol{\tau}\right]\right]\right\}\times \\
& \left[\int \exp\left\{-\frac{1}{2}\left[(\boldsymbol{\mu}-\tilde{\boldsymbol{\mu}}^\mu)^T\tilde{\boldsymbol{\tau}}^\mu(\boldsymbol{\mu}-\tilde{\boldsymbol{\mu}}^\mu)\right]\right\}d\boldsymbol{\mu}\right]d\boldsymbol{\Sigma} \qquad \text{(S46)}
\end{aligned}
$$

where the effective covariance has the form,

$$
\tilde{\boldsymbol{\Sigma}} = m^0(\boldsymbol{y}-\tilde{\boldsymbol{\mu}}^y)(\boldsymbol{y}-\tilde{\boldsymbol{\mu}}^y)^T + m^1(\boldsymbol{\mu}_0-\bar{\boldsymbol{x}})(\boldsymbol{\mu}_0-\bar{\boldsymbol{x}})^T + n\boldsymbol{S}_x^2 + \boldsymbol{\Sigma}_0 \qquad \text{(S47)}
$$

The mean parameter in (S46) only appears in the term in brackets, so we can integrate it in a straightforward manner as a multivariate normal integral:

$$
\begin{aligned}
p(\boldsymbol{y}|\{\boldsymbol{x}\}) \propto & \int \det[\boldsymbol{\tau}]^{(n+\nu_0+p+3)/2}\exp\left\{-\frac{1}{2}\left[\mathrm{Tr}\left[\tilde{\boldsymbol{\Sigma}}\boldsymbol{\tau}\right]\right]\right\}\det[\tilde{\boldsymbol{\tau}}]^{-1/2}d\boldsymbol{\Sigma} \\
& \propto \int \det[\boldsymbol{\tau}]^{(n+\nu_0+p+2)/2}\exp\left\{-\frac{1}{2}\left[\mathrm{Tr}\left[\tilde{\boldsymbol{\Sigma}}\boldsymbol{\tau}\right]\right]\right\}d\boldsymbol{\Sigma} \qquad \text{(S48)}
\end{aligned}
$$

where $\det[\tilde{\boldsymbol{\tau}}^\mu] \propto \det[\boldsymbol{\tau}]$. The last integral is an inverse Wishart that we can integrate directly, leading to

$$
p(\boldsymbol{y}|\{\boldsymbol{x}\}) \propto \det[\tilde{\boldsymbol{\Sigma}}]^{-(n+\nu_0+1)/2} \qquad \text{(S49)}
$$

We can now reorganize the effective covariance. If we define the matrix

$$
\tilde{\boldsymbol{\Sigma}}^y = \frac{1}{m^0}\left(m^1(\boldsymbol{\mu}_0-\bar{\boldsymbol{x}})(\boldsymbol{\mu}_0-\bar{\boldsymbol{x}})^T + n\boldsymbol{S}_x^2 + \boldsymbol{\Sigma}_0\right), \qquad \text{(S50)}
$$

we can rewrite the effective covariance $\tilde{\boldsymbol{\Sigma}}$ as

$$
\tilde{\boldsymbol{\Sigma}} = \left((\boldsymbol{y}-\tilde{\boldsymbol{\mu}}^y)(\boldsymbol{y}-\tilde{\boldsymbol{\mu}}^y)^T\tilde{\boldsymbol{\tau}}^y + \mathbb{I}\right)m_0\tilde{\boldsymbol{\Sigma}}^y, \qquad \text{(S51)}
$$

Insert this expression in equation (S49) we obtain

$$
\begin{aligned}
p(\boldsymbol{y}|\{\boldsymbol{x}\}) & \propto \det[\left((\boldsymbol{y}-\tilde{\boldsymbol{\mu}}^y)(\boldsymbol{y}-\tilde{\boldsymbol{\mu}}^y)^T\tilde{\boldsymbol{\tau}}^y + \mathbb{I}\right)n_0\tilde{\boldsymbol{\Sigma}}^y] \\
& \propto \det[(\boldsymbol{y}-\tilde{\boldsymbol{\mu}}^y)(\boldsymbol{y}-\tilde{\boldsymbol{\mu}}^y)^T\tilde{\boldsymbol{\tau}}^y + \mathbb{I}]^{-(n+\nu_0+1)/2} = |1+(\boldsymbol{y}-\tilde{\boldsymbol{\mu}}^y)^T\tilde{\boldsymbol{\tau}}^y(\boldsymbol{y}-\tilde{\boldsymbol{\mu}}^y)|^{-(n+\nu_0+1)/2} \\
& = |1+\frac{1}{\tilde{\nu}}(\boldsymbol{y}-\tilde{\boldsymbol{\mu}}^y)^T(\tilde{\nu}\tilde{\boldsymbol{\tau}}^y)(\boldsymbol{y}-\tilde{\boldsymbol{\mu}}^y)|^{-(\tilde{\nu}+p)/2} = t_{\tilde{\nu}}(\boldsymbol{y};\tilde{\boldsymbol{\mu}}^y,\tilde{\boldsymbol{\Sigma}}^y/\tilde{\nu}) \qquad \text{(S52)}
\end{aligned}
$$

which is a multivariate T-distribution with

$$m^0 = \frac{n+\kappa_0}{n+\kappa_0+1}, \quad m^1 = \frac{\frac{n\kappa_0}{n+\kappa_0}+n\kappa_0}{n+\kappa_0+1}, \quad \tilde{\nu} = n+\nu_0+1-p \tag{S53}$$

$$\tilde{\boldsymbol{\mu}}^y = \frac{(n\bar{\boldsymbol{x}}+\kappa_0\boldsymbol{\mu}_0)}{n+\kappa_0} \tag{S54}$$

$$\tilde{\boldsymbol{\Sigma}}^y = \frac{1}{m^0}\left(m^1(\boldsymbol{\mu}_0-\bar{\boldsymbol{x}})(\boldsymbol{\mu}_0-\bar{\boldsymbol{x}})^T + n\boldsymbol{S}_x^2 + \boldsymbol{\Sigma}_0\right) \tag{S55}$$

#### S3.2 Finite mixture distributions

We derive in this section the statistics relevant to mixture models.

##### S3.2.1 Likelihood

The likelihood of a finite mixture model with $K$ components has the form

$$p(\{\boldsymbol{x}\}|\{\boldsymbol{\phi}\},\boldsymbol{\omega}) = \prod_i \left(\sum_j^K \omega_j p(\boldsymbol{x}_i|\boldsymbol{\phi}_j)\right), \tag{S56}$$

where the vector sets $\{\boldsymbol{x}\} = \{\boldsymbol{x}_i : i = 1\dots N\}$ and $\{\boldsymbol{\phi}\} = \{\boldsymbol{\phi}_j : j = 1\dots K\}$ run over the number of samples (cells) $N$ and the number of mixture components $K$, respectively (in what follows we use the subindices $i$ and $j$ with those two distinct meanings)[1]. The set of parameters $\boldsymbol{\omega}$ are called the weights of the mixture model, and $p(\{\boldsymbol{x}\}|\boldsymbol{\phi}_j)$ are the base distributions. We can extend this model to introduce a set of hidden indicator variables $\{\boldsymbol{z}\}$, defined as

$$z_{ij} = \delta_{jk_i} \tag{S57}$$

for some $k_i \in \{1,...,K\}$, and where $\delta_{ij}$ is the Kronecker delta. This variable basically tells from which distribution $p(\{\boldsymbol{x}\}|\boldsymbol{\phi}_i)$ the variable came from. Using this set of hidden variables, our model (S56) can be rewritten as

$$p(\{\boldsymbol{x}\},\{\boldsymbol{z}\}|\{\boldsymbol{\phi}\},\boldsymbol{\omega}) = \prod_i^N \prod_j^K p(\boldsymbol{x}_i|\boldsymbol{\phi}_j)^{z_{ij}}\omega_j^{z_{ij}} \tag{S58}$$

It is straightforward to see that, if we take the marginal distribution over the hidden variables, we recover the original distribution:

$$p(\{\boldsymbol{x}\}|\{\boldsymbol{\phi}\},\boldsymbol{\omega}) = \prod_i^N \left(\sum_j^K p(\boldsymbol{x}_i, z_{ij}=1|\boldsymbol{\phi}_j,\boldsymbol{\omega})\right) = \prod_i^N \left(\sum_j^K p(\boldsymbol{x}_i|\boldsymbol{\phi}_j)\omega_j\right) \tag{S59}$$

[1] We remind the reader that the dimension of the vectors $\boldsymbol{x}_i$ and $\boldsymbol{\phi}_j$ is equal to the number of measurement channels.

where only when the indicator variable is one of the corresponding term survives.

The use of indicator variables makes it possible to compute analytically the posterior distribution of the mixture model, as the base distributions are now in product form. The hidden indicator variables are not known, thus we will have to sample from them as well.

###### S3.2.2 Conjugate prior

A conjugate prior for the mixture model is

$$p(\boldsymbol{\omega}|\boldsymbol{\alpha}) = \text{Dirichlet}(\boldsymbol{\omega}|\boldsymbol{\alpha}) \tag{S60}$$

where the Dirichlet distribution has the form

$$\text{Dirichlet}(\boldsymbol{\omega}|\boldsymbol{\alpha}) \propto \prod_{i}^{K} \omega_i^{\alpha_i - 1} \tag{S61}$$

The hyperprior parameters $\alpha$ are usually set to be symmetrical and to scale with the number of mixture components:

$$\alpha_i = \alpha/K \quad \forall i \tag{S62}$$

In this way, the prior distribution only depends on one hyperparameter $\alpha$ that indicates the strength from the uniform weights.

###### S3.2.3 Posterior distribution

Putting together the likelihood and the prior distribution, the posterior of the mixture model is

$$p(\{\boldsymbol{\phi}\}, \boldsymbol{\omega}|\{\boldsymbol{x}\}, \{\boldsymbol{z}\}) \propto \prod_{j}^{K} \left( \prod_{i} p(\boldsymbol{x}_i|\boldsymbol{\phi}_j)^{z_{ij}} \omega_j^{z_{ij}} \right) \omega_j^{\alpha/K-1} p(\boldsymbol{\phi}_j^0), \tag{S63}$$

where the last term is the set of priors for each base distribution.

###### S3.2.4 Conditional distribution: weights

Taking the terms from the posterior that involve the weights:

$$p(\boldsymbol{\omega}|\{\boldsymbol{z}\}) \propto \prod_{j}^{K} \left( \prod_{i} \omega_j^{z_{ij}} \right) \omega_j^{\alpha/K-1} = \text{Dirichlet}(\boldsymbol{\omega}|\tilde{\boldsymbol{n}}) \tag{S64}$$

where $\tilde{n}_j = \sum_i z_{ij} + \alpha$.

###### S3.2.5 Conditional distribution: indicator variables

As we already mentioned, the indicator variables are not known, so we have to sample from them too. As we are considering identically independent samples, we can obtain the conditional distribution from each indicator variable independently as

$$p(\boldsymbol{c}_i|\boldsymbol{\omega}, \{\boldsymbol{\phi}\}, \boldsymbol{x}_i) \propto \prod_j^K p(\boldsymbol{x}_i|\boldsymbol{\phi}_j)^{z_{ij}} \omega_j^{z_{ij}} = \text{Multinomial}(\boldsymbol{z}_i; 1, \{p(\boldsymbol{x}_i|\boldsymbol{\phi}_j)\omega_j\}_j) \tag{S65}$$

It is worth noting that the indicator will be sampled from a particular base distribution for the weights of that base, but also by how well that sample lies inside the base distribution.

###### S3.2.6 Conditional distribution: base distribution parameters

Finally, because the base distributions are in product form due to the introduction of the indicator variables, the parameters of each base can be computed independently as

$$p(\boldsymbol{\phi}_j|\{\boldsymbol{x}\}, \{\boldsymbol{z}\}) \propto p(\{\boldsymbol{x}\}_j|\boldsymbol{\phi}_j)p(\boldsymbol{\phi}_j^0) \tag{S66}$$

where $\{\boldsymbol{x}\}_j$ represents the subsample of cells whose indicator variable belongs to the corresponding mixture component.

##### S3.3 Infinite mixture distributions

In this section we define basic results from infinite Dirichlet processes (for an insightful tutorial see [2]) that allow us to consider infinite mixtures.

###### S3.3.1 Taking the infinite limit

In order to take the limit to infinite clusters, we need to remove the dependence from the number of clusters in a mixture model. For that, we need to remove the dependence on the weights of our probability distribution. Consider for the moment a basis distribution that is uniform in space. The joint probability distribution conditioned on the priors would be

$$p(\{\boldsymbol{z}\}, \boldsymbol{\omega}|\alpha) = p(\{\boldsymbol{z}\}|\boldsymbol{\omega})p(\boldsymbol{\omega}|\alpha) \tag{S67}$$

where the first term in the right-hand side is the likelihood of the indicator variables as in (S58), also given in (S65), which is a multinomial distribution. The second term is the prior distribution of the mixture distribution (S61), which is a Dirichlet distribution. From this expression, we can calculate the marginal distribution of the indicator variables conditioned to the prior parameter:

$$
\begin{aligned}
p(\{\boldsymbol{z}\}|\boldsymbol{\alpha}) &= \int p(\{\boldsymbol{z}\}|\boldsymbol{\omega})p(\boldsymbol{\omega}|\boldsymbol{\alpha})d\boldsymbol{\omega} = \int \prod_i \left(\prod_j \omega_j^{z_{ij}}\right) \frac{\Gamma(\alpha)}{\Gamma(\alpha/K)^K} \prod_j \omega_j^{\alpha/K-1} d\boldsymbol{\omega} \\
&= \frac{\Gamma(\alpha)}{\Gamma(\alpha/K)^K} \int \prod_j \omega_j^{n_j+\alpha/K-1} d\boldsymbol{\omega} = \frac{\Gamma(\alpha)}{\Gamma(\alpha/K)^K} \frac{\prod_j \Gamma(n_j+\alpha/K)}{\Gamma(n+\alpha/K)}
\end{aligned} \tag{S68}
$$

where the third equality makes use of the statistic $n_j = \sum_i z_{ij}$. The integral can be identified as a multinomial distribution without the scaling factor.

The expression above still contains explicitly the dependence on the number of components in the mixture $K$, and thus its limit $K \to \infty$ cannot be computed in a straightforward manner. To have a more amenable expression, let us consider that we take out the sample $l$ and reassign it to a new cluster. The probability of this sample to be assigned to any of the clusters, conditioned on all the other indicator variables, is

$$
\begin{aligned}
p(z_{lk}=1|\{\boldsymbol{z}\}_{\neg l},\alpha) &= \frac{p(\{\boldsymbol{z}\}_{\neg l}, z_{lk}=1|\alpha)}{p(p(\{\boldsymbol{z}\}_{\neg l}|\alpha)} = \frac{\frac{\Gamma(\alpha)}{\Gamma(\alpha/K)^K}\frac{\prod_j \Gamma(n_{j|l}+\delta_{jk}+\alpha/K)}{\Gamma(n+\alpha/K)}}{\frac{\Gamma(\alpha)}{\Gamma(\alpha/K)^K}\frac{\prod_j \Gamma(n_{j|l}+\alpha/K)}{\Gamma(n+\alpha/K-1)}} \\
&= \frac{\Gamma(n+\alpha/K-1)}{\Gamma(n+\alpha/K)} \prod_j \frac{\Gamma(n_{j|l}+\delta_{jk}+\alpha/K)}{\Gamma(n_{j|l}+\alpha/K)} = \frac{n_{k|l}+\alpha/K}{n+\alpha-1},
\end{aligned} \tag{S69}
$$

where have made explicit the range of cells over which the samples $\{\boldsymbol{z}\}$ are taken in each case. To that end, we define $\{\boldsymbol{z}\}_{\neg l} \equiv \{\boldsymbol{z}\}_{i\in(1,\ldots,l-1,l+1,\ldots n)}$ and $n_{\neg l} \equiv \sum_{i\in(1,\ldots,l-1,l+1,\ldots n)} z_{ij}$ to refer to the subsets that contain all elements except $l$. With this expression, it is straightforward to take the limit to infinite components:

$$
p_\infty(z_{lk}=1|\{\boldsymbol{z}\}_{\neg l},\alpha) = \lim_{K\to\infty} p(z_{lk}=1|\{\boldsymbol{z}\}_{\neg l},\alpha) = \frac{n_{\neg l}}{n+\alpha-1} \tag{S70}
$$

We now have the probability that a sample is assigned to a base that has other indicator variables. Let us now consider for the moment that we have $K$ bases with assigned samples. The probability that a sample is assigned to a new base will be:

$$
\begin{aligned}
p_\infty(\boldsymbol{z}_{l,K+1}=1|\{\boldsymbol{z}\}_{\neg l},\alpha) &= 1 - \sum_{j=1}^{K} p_\infty(z_{lk}=1|\{\boldsymbol{z}\}_{\neg l},\alpha) \\
&= 1 - \sum_{j=1}^{K} \frac{n_{k|l}}{n+\alpha-1} = 1 - \frac{n-1}{n+\alpha-1} = \frac{\alpha}{n+\alpha-1}
\end{aligned} \tag{S71}
$$

There is a non-zero probability that the cell will be assigned to a new base that was not populated before, and the probability of populating this new cluster will depend on the hyperparameter $\alpha$.

##### S3.3.2 Adding a non-uniform basis

The results above have been derived considering a uniform basis distribution. In the most general case, the basis will be non-uniform. From Eq. (S58) it is very easy to see that if we had a non-uniform distribution, the corresponding term will drop out of the integral (S68). Proceeding in the same way as before, all the terms in (S69) will cancel out, except $p(x_l|\boldsymbol{\phi}_k)$.

The probabilities of a new assignation will be

$$p_\infty(z_{lk} = 1|\{\boldsymbol{z}\}_{\neg l}, \{\boldsymbol{\phi}\}, \alpha) \propto \frac{n_{\neg l}}{n+\alpha-1} p(\boldsymbol{x}_i|\boldsymbol{\phi}_k) \tag{S72}$$

for a basis with a populated sample and

$$p_\infty(z_{l,K+1} = 1|\{\boldsymbol{z}\}_{\neg l}, \{\boldsymbol{\phi}\}, \alpha) \propto \frac{\alpha}{n+\alpha-1} p(\boldsymbol{x}_i|\boldsymbol{\phi}^0) \tag{S73}$$

for the creation of a new basis.

##### S3.3.3 Using the predictive posterior distribution

Similar way to the approach used in Sec. S3.1.7, we can directly predict the new outcomes in the case of an infinite mixture as a function of already observed data.

The probabilities of a new assignation will be

$$p_\infty(z_{lk} = 1|\{\boldsymbol{z}\}_{\neg l}, \{\boldsymbol{\phi}\}, \alpha) \propto \frac{n_{\neg l}}{n+\alpha-1} p(\boldsymbol{x}_i|\{\boldsymbol{x}\}_k) \tag{S74}$$

for a basis with a populated sample and

$$p_\infty(z_{l,K+1} = 1|\{\boldsymbol{z}\}_{\neg l}, \{\boldsymbol{\phi}\}, \alpha) \propto \frac{\alpha}{n+\alpha-1} p(\boldsymbol{x}_i|\boldsymbol{\phi}^0) \tag{S75}$$

for a new basis.

#### S3.4 Modifications of the convolution distribution

Introducing indicator variables as indicated in S3.2.1, the convolution of two multivariate mixture distributions like the one described by (S11) takes the form

$$p(\{\boldsymbol{x}\}, \{\boldsymbol{z}\}|\{\boldsymbol{\phi}^\xi\}, \boldsymbol{\omega}^\xi, \{\boldsymbol{\phi}^T\}, \boldsymbol{\omega}^T) = \prod_i^N \prod_j^{K^\xi} \prod_k^{K^T} \mathcal{N}(\boldsymbol{x}_i|\boldsymbol{\mu}_j^\xi + \boldsymbol{\mu}_k^T, \boldsymbol{\Sigma}_j^\xi + \boldsymbol{\Sigma}_k^T)^{z_{ijk}} (\omega_j^\xi \omega_k^T)^{z_{ijk}} \tag{S76}$$

where the indicator variable is now 1 if the sample $i$ belongs to the noise base $j$ and target base $k$. Considering the approximation described in S2.1, the only sampling parameters that we have to go over will be $\{\boldsymbol{\phi}^T\}$, the weights $\boldsymbol{\omega}^T$ and the indicator variables $\boldsymbol{z}$. The main challenge is that there is no close form to group all the terms involving $\Sigma^T$ in single effective distributions as the ones derived in S3.1. In order to get a tractable expression, we would like to transform the convoluted covariances in such a way that the following expression follows:

$$\boldsymbol{M}(\boldsymbol{\Sigma}_j^\xi + \boldsymbol{\Sigma}_k^T)^{-1} = (\boldsymbol{M} + \boldsymbol{A})\boldsymbol{\Sigma}_k^{T^{-1}} \tag{S77}$$

for given matrices $\boldsymbol{A}$, $\boldsymbol{M}$. Isolating, $\boldsymbol{A}$ we obtain that

$$\boldsymbol{A} = -\boldsymbol{M}(\boldsymbol{\Sigma}_j^\xi + \boldsymbol{\Sigma}_k^T)^{-1}\boldsymbol{\Sigma}_j^\xi \tag{S78}$$

Now, in most practical cases we can consider that the convoluted covariance will be close to the expected covariance matrix, and we can approximate the expression above by

$$\boldsymbol{A} \sim -\boldsymbol{M}\hat{\boldsymbol{\tau}}_{jk}\boldsymbol{\Sigma}_j^\xi, \tag{S79}$$

where the effective expected covariance is

$$\hat{\boldsymbol{\Sigma}}_{jk} = n_{jk}\boldsymbol{S}_{jk}^2 + \kappa_0(\bar{\boldsymbol{x}}_{jk} - \boldsymbol{\mu}_0)(\bar{\boldsymbol{x}}_{jk} - \boldsymbol{\mu}_0)^T + \boldsymbol{\Sigma}_0 \tag{S80}$$

All the results derived in the preceding sections take into account this approximation:

$$\boldsymbol{\mu} \rightarrow \boldsymbol{\mu}^T + \boldsymbol{\mu}^\xi \tag{S81}$$

$$\boldsymbol{\Sigma} \rightarrow \boldsymbol{\Sigma}^T + \boldsymbol{\Sigma}^\xi \tag{S82}$$

$$\boldsymbol{\tau}^T \rightarrow (\mathbb{I} - \hat{\boldsymbol{\tau}}_{jk}\boldsymbol{\Sigma}_j^\xi)\boldsymbol{\tau}^T \tag{S83}$$

$$z_{ij} \rightarrow z_{ijk} \tag{S84}$$

The weights of the target distribution (S3.2.4) will be computed using the sum over the samples for all the noise samples

$$n_k^T = \sum ijz_{ijk} \tag{S85}$$

Sampling from the mean and covariance parameters in the distribution needs, however, a closer look.

##### S3.4.1 Prior distribution

In order for the prior to be conjugated, we have to scale the prior distribution in terms of the prior of the noise distribution.

$$p(\boldsymbol{\mu}|\tilde{\boldsymbol{\Sigma}}) = \mathcal{N}(\boldsymbol{\mu}; \boldsymbol{\mu}_0, \kappa_0 \tilde{\boldsymbol{\Sigma}}) \tag{S86}$$

$$p(\tilde{\boldsymbol{\Sigma}}) = \mathcal{W}^{-1}(\tilde{\boldsymbol{\Sigma}}; \boldsymbol{\Sigma}_0, \boldsymbol{\Sigma}^{\xi}, \nu_0) \tag{S87}$$

where the effective covariance is

$$\tilde{\boldsymbol{\Sigma}} = \boldsymbol{\Sigma}^T + \boldsymbol{\Sigma}^{\xi} \tag{S88}$$

##### S3.4.2 Conditional distribution: covariance

If we focus the analysis in a single set of target parameters $\boldsymbol{\phi}_k$, the posterior probability of the variance and the mean has the form

$$p({\boldsymbol{\phi}^T}_k|\{\boldsymbol{\phi}^{\xi}\}, \{\boldsymbol{x}\}, \{\boldsymbol{z}\}) \propto \prod_i^N \prod_j^{K_\xi} \mathcal{N}(\boldsymbol{x}_i|\boldsymbol{\mu}_j^{\xi} + \boldsymbol{\mu}_k^T, \boldsymbol{\Sigma}_j^{\xi} + \boldsymbol{\Sigma}_k^T)^{z_{ijk}} p(\boldsymbol{\mu}|\boldsymbol{\Sigma}_j^{\xi} + \boldsymbol{\Sigma}_k^T) p(\boldsymbol{\Sigma}_j^{\xi} + \boldsymbol{\Sigma}_k^T) \tag{S89}$$

In contrast with the case without convolution, the normal distributions cannot be grouped together in a simple distribution that can be sampled by standard procedures. We can group the data coming from a specific dataset,

$$p({\boldsymbol{\phi}^T}_k|\{\boldsymbol{\phi}^{\xi}\}, \{\boldsymbol{x}\}, \{\boldsymbol{z}\}) \propto \det\left(\boldsymbol{\Sigma}_j^{\xi} + \boldsymbol{\Sigma}_k^T\right)^{-(n_{jk}+\nu_0+1)/2} \exp\left\{-\frac{1}{2}\left[\tilde{\boldsymbol{\Sigma}}_{jk}(\boldsymbol{\Sigma}_j^{\xi} + \boldsymbol{\Sigma}_k^T)^{-1}\right]\right\} \tag{S90}$$

where the variance is

$$\tilde{\boldsymbol{\Sigma}}_{jk} = n_{jk}\boldsymbol{S}_{jk}^2 + n_{jk}(\bar{\boldsymbol{x}}_{jk} - \boldsymbol{\mu}_j^{\xi} - \boldsymbol{\mu}_k^T)(\bar{\boldsymbol{x}} - \boldsymbol{\mu}_j^{\xi} - \boldsymbol{\mu}_k^T)^T + \kappa_0(\boldsymbol{\mu}_k^T - \boldsymbol{\mu}_0)(\boldsymbol{\mu}_k^T - \boldsymbol{\mu}_0)^T + \boldsymbol{\Sigma}_0 \tag{S91}$$

Within the proposed approximation, we can now group all the different distributions:

$$\tilde{\boldsymbol{\Sigma}}_k = \sum_j \tilde{\boldsymbol{\Sigma}}_{jk}(\mathbb{I} - \hat{\boldsymbol{\tau}}_{jk}\boldsymbol{\Sigma}_j^{\xi}) \tag{S92}$$

and we can sample new covariance matrices for the target with the inverse Wishart distribution.

$$\boldsymbol{\Sigma}_k^T \sim \mathcal{W}^{-1}(\tilde{\boldsymbol{\Sigma}}_k, n_k + \nu_0 + 1) \tag{S93}$$

##### S3.4.3 Conditional distribution: mean

The mean distribution can be easily computed, regrouping all the terms in the following summary statistics:

$$\tilde{\boldsymbol{\tau}}_k = \sum_j (n_{jk} + \kappa_0)(\boldsymbol{\Sigma}_j^{\xi} + \boldsymbol{\Sigma}_k^T)^{-1}$$
$$\tilde{\boldsymbol{\mu}}_k = \tilde{\boldsymbol{\Sigma}}_j \left( \sum_j (\boldsymbol{\Sigma}_j^{\xi} + \boldsymbol{\Sigma}_k^T)^{-1} (n_{jk}\bar{\boldsymbol{x}}_{jk} + \kappa_0 \boldsymbol{\mu}_0) \right),$$

and sampling from a multivariate normal,

$$\boldsymbol{\mu}_k^T \sim \mathcal{N}(\tilde{\boldsymbol{\mu}}_k, \tilde{\boldsymbol{\tau}}_k) \quad \text{(S94)}$$

### S4 Hyperparameter selection

Our approach the has five hyperparameters:

- $\alpha$: Determines how close we are from a uniform distribution of weights (finite mixtures), or the potential of generating a new basis (infinite mixture).
- $\boldsymbol{\mu}_0$: The center of the prior normal distribution $\mu_0$.
- $\kappa_0$: The confidence we have of being close to $\mu_0$.
- $\boldsymbol{\Sigma}_0$: The covariance of the prior normal distribution.
- $\nu_0$: The confidence we have of being close to $\sigma_0$.

Flow cytometry applications have in general have large datasets, making the approach quite insensitive to the choice of $\alpha$, $\kappa_0$ and $\nu_0$, as the statistics will dominate the model. The choice of mean and covariance matrix, however, depend on the data distribution. Appropriate estimators of these parameters are the mean and the covariance matrix of the whole dataset. This imposes a soft-informative prior over the region where the density should lie. In cases where the distribution has fat tails, the density prior can be very flat because of the outliers, making it difficult for the algorithm to find the correct distribution as the prior spreads over the basis components. In these cases, narrower prior covariance matrices can help to fit the model correctly.

#### S5 Gibbs sampling algorithms

In this section, we describe the algorithms to sample the noise distribution according to the approximation described in S2.1.

##### S5.1 Efficient updating of summary statistics

Once we have computed the mean or the variance over a set of samples $\{\boldsymbol{x}\}$ and we add or remove a single sample, it is possible to update the statistic without having to recompute the metric fully over the new set, which in general will be more time-consuming as we will have to go over a sum over all the size of the set.

If we remove a sample:

$$n_j^{rem} = n_j - 1 \tag{S95}$$

$$\bar{\boldsymbol{x}}_j^{rem} = \frac{n_j \bar{\boldsymbol{x}}_j - \boldsymbol{x}_i}{n_j^{rem}} \tag{S96}$$

$$\boldsymbol{S}_j^{2\,rem} = \frac{1}{n_j^{rem}}(n_j \boldsymbol{S}_j^2 + n_j \bar{\boldsymbol{x}}_j \bar{\boldsymbol{x}}_j^T - \boldsymbol{x}_i \boldsymbol{x}_i^T) - \bar{\boldsymbol{x}}_j^{rem} \bar{\boldsymbol{x}}_{j,rem}^T \tag{S97}$$

On the other hand, if we add a sample:

$$n_j^{add} = n_j + 1 \tag{S98}$$

$$\bar{\boldsymbol{x}}_j^{add} = \frac{n_j \bar{\boldsymbol{x}}_j + \boldsymbol{x}_i}{n_j^{add}} \tag{S99}$$

$$\boldsymbol{S}_j^{2\,add} = \frac{1}{n_j^{add}}(n_j \boldsymbol{S}_j^2 + n_j \bar{\boldsymbol{x}}_j \bar{\boldsymbol{x}}_j^T + \boldsymbol{x}_i \boldsymbol{x}_i^T) - \bar{\boldsymbol{x}}_j^{new} \bar{\boldsymbol{x}}_{j,add}^T \tag{S100}$$

##### S5.2 Finite normal mixture distribution

The procedure in this case is as follows:

0 Initialize the parameters.

0.1 Initialize the indicator variables $\{\boldsymbol{z}\}$: Assign to each cell one of the $K$ normal distributions using any initialization procedure (random assignment, k-means...).

0.2 Compute the statistics of each base:

$$n_j = \sum_i z_{ij} \qquad \bar{\boldsymbol{x}}_j = \frac{1}{n_j}\sum_i z_{ij}\boldsymbol{x}_i \qquad \boldsymbol{S}_j^2 = \frac{1}{n_j}\sum_i z_{ij}\boldsymbol{x}_i\boldsymbol{x}_i^T - \bar{\boldsymbol{x}}_j\bar{\boldsymbol{x}}_j^T$$

0.3 Initialize parameters of each base:

$$\boldsymbol{\mu}_j = \bar{\boldsymbol{x}}_j \qquad \boldsymbol{\Sigma}_j = \boldsymbol{S}_j^2 \qquad \omega_j = \frac{1}{n}\sum_i z_{ij}$$

where $n = \sum_j n_j$ is the size of the dataset.

1 Sampling. For $N$ iterations do

1.1 Sample indicator variables as in S3.2.5. For each sample $i$:

1.1.1 Compute weights: $w_j = \omega_j p(\boldsymbol{x}_i|\boldsymbol{\phi}_j)$.

1.1.2 Sample new indicator: $\boldsymbol{z} \sim \text{Multinomial}(1, \boldsymbol{w})$.

1.2 Sample weights as from S3.2.4. For each base $j \in 1, ...K$ do

1.2.1 Compute $\tilde{\boldsymbol{n}}$: $\tilde{n}_j = \sum_i z_{ij} + \alpha/K$

1.2.2 Sample new weights: $\boldsymbol{\omega} \sim \text{Dirichlet}(\tilde{\boldsymbol{n}})$

1.3 Sample variances and mean. For each base $j \in 1, ...K$ do

1.3.1 Compute summary statistics:

$$n_j = \sum_i z_{ij} \qquad \bar{\boldsymbol{x}}_j = \frac{1}{n_j}\sum_i z_{ij}\boldsymbol{x}_i \qquad \boldsymbol{S}_j^2 = \frac{1}{n_j}\sum_i z_{ij}\boldsymbol{x}_i\boldsymbol{x}_i^T - \bar{\boldsymbol{x}}_j\bar{\boldsymbol{x}}_j^T$$

1.3.2 Sample new covariance S3.1.5:

$$\begin{aligned}
\tilde{n}_j &= n_j + \nu_0 + 1 \\
\tilde{\boldsymbol{\Sigma}}_j &= n_j\boldsymbol{S}_j^2 + n_j(\bar{\boldsymbol{x}} - \boldsymbol{\mu}_j)(\bar{\boldsymbol{x}} - \boldsymbol{\mu}_j)^T + \kappa_0(\boldsymbol{\mu}_j - \boldsymbol{\mu}_0)(\boldsymbol{\mu}_j - \boldsymbol{\mu}_0)^T + \boldsymbol{\Sigma}_0 \\
\boldsymbol{\Sigma}_j &\sim \mathcal{W}^{-1}(\tilde{n}_j, \tilde{\boldsymbol{\Sigma}}_j)
\end{aligned}$$

1.3.2 Sample new mean S3.1.4:

$$\begin{aligned}
\tilde{\boldsymbol{\mu}} &= \frac{n_j\bar{\boldsymbol{x}}_j + \kappa_0\boldsymbol{\mu}_0}{n_j + \kappa_0} \\
\tilde{\boldsymbol{\tau}} &= (n_j + \kappa_0)\boldsymbol{\tau}_j \\
\boldsymbol{\mu}_j &\sim \mathcal{N}(\tilde{\boldsymbol{\mu}}, \tilde{\boldsymbol{\Sigma}})
\end{aligned}$$

#### S5.3 Infinite normal mixture distribution

The approach in this case has the following steps:

0 Initialize the parameters.

0.1 Initialize the indicator variables $\{\boldsymbol{z}\}$: Assign to each cell one of the $K$ normal distributions using any initialization procedure (random assignment, k-means...).

0.2 Compute the statistics of each base:

$$n_j = \sum_i z_{ij} \qquad \bar{\boldsymbol{x}}_j = \frac{1}{n_j}\sum_i z_{ij}\boldsymbol{x}_i \qquad \boldsymbol{S}_j^2 = \frac{1}{n_j}\sum_i z_{ij}\boldsymbol{x}_i\boldsymbol{x}_i^T - \bar{\boldsymbol{x}}_j\bar{\boldsymbol{x}}_j^T$$

0.3 Initialize parameters of each base:

$$\boldsymbol{\mu}_j = \bar{\boldsymbol{x}}_j \qquad \boldsymbol{\Sigma}_j = \boldsymbol{S}_j^2 \qquad \omega_j = \frac{1}{n}\sum_i z_{ij}$$

where $n = \sum_j n_j$ is the size of the dataset.

1 Sampling. For $N$ iterations, do

1.1 Reassign samples. For each sample $i \in \{1, ..., n\}$

1.1.1 Remove sample $i$. Consider sample $z_{ij} = 1$.

If $n_j = 1$ (the only sample assigned to that base distribution), remove the distribution from the active basis.

Otherwise recompute the summary statistics for base distribution $j$ removing one sample as described in S5.1.

1.1.2 Compute weights (not-normalized) for reassigning the sample as described in sections S3.3.2 and S3.1.7.

For the active bases $j \in \{1, ...K\}$ .

$$m_j^0 = \frac{n_j + \kappa_0}{n_j + \kappa_0 + 1} \qquad m_j^1 = \frac{\frac{n_j\kappa_0}{n_j+\kappa_0} + n_j\kappa_0}{n_j + \kappa_0 + 1}$$

$$\tilde{\boldsymbol{\mu}}_j^y = \frac{(n_j\bar{\boldsymbol{x}} + \kappa_0\boldsymbol{\mu}_0)}{n_j + \kappa_0} \qquad \tilde{\boldsymbol{\Sigma}}_j^y = \frac{1}{m^0}\left(m^1(\boldsymbol{\mu}_0 - \bar{\boldsymbol{x}})(\boldsymbol{\mu}_0 - \bar{\boldsymbol{x}})^T + n_j\boldsymbol{S}_x^2 + \boldsymbol{\Sigma}_0\right)$$

$$w_j = \frac{n_j}{n + \alpha - 1}t_{\tilde{\nu}}(\boldsymbol{y}; \tilde{\boldsymbol{\mu}}^y, \tilde{\boldsymbol{\Sigma}}^y/\tilde{\nu})$$

and for creating a new basis,

$$w_{K+1} = \frac{\alpha}{n + \alpha - 1}\mathcal{N}(\boldsymbol{x}_i|\boldsymbol{\mu}_0^y, \boldsymbol{\Sigma}_0^y)$$

1.1.3 Sample new indicator: $\boldsymbol{z}_i^{new} \sim \text{Multinomial}(1, \boldsymbol{w})$.

1.1.4 Update statistics.

If $z_{iK+1} = 1$ create a new basis distribution and assign new statistics to it,

$$n_{K+1} = 1 \qquad \bar{\boldsymbol{x}}_{K+1} = \boldsymbol{x}_i \qquad \boldsymbol{S}_{K+1}^2 = 0$$

else, update the statistics adding a term as described in S5.1.

1.2 Sample basis parameters. For each basis $j \in \{1, ..., K\}$:

1.2.1 Sample the weights S3.2.4:

$$\tilde{n}_j = n_j + \alpha \qquad \boldsymbol{\omega} \sim \text{Dirichlet}(\tilde{\boldsymbol{n}})$$

1.2.2 Sample the covariance matrix S3.1.6:

$$\tilde{n}_j = n_j + \nu_0 + 1 \qquad \tilde{\boldsymbol{\Sigma}}_j = n\boldsymbol{S}_j^2 + \frac{\kappa_0 n_j}{\kappa_0 + n_j}(\bar{\boldsymbol{x}}_j - \boldsymbol{\mu}_0)(\bar{\boldsymbol{x}}_j - \boldsymbol{\mu}_0)^T + \boldsymbol{\Sigma}_0$$

$$\boldsymbol{\Sigma}_j \sim \mathcal{W}^{-1}(\tilde{\boldsymbol{\Sigma}}_j, \tilde{n}_j)$$

1.2.3 Sample the mean S3.1.4:

$$\tilde{\boldsymbol{\mu}}_j = \frac{n_j \bar{\boldsymbol{x}}_j + \kappa_0 \boldsymbol{\mu}_0}{n_j + \kappa_0} \qquad \tilde{\boldsymbol{\Sigma}}_j = \boldsymbol{\Sigma}_j/(n_j + \kappa_0)$$

$$\boldsymbol{\mu}_j \sim \mathcal{N}(\tilde{\boldsymbol{\mu}}_j, \tilde{\boldsymbol{\Sigma}}_j)$$

##### S5.4 Modifications to the convoluted distribution

The preceding algorithms are the sampling algorithms for non-convoluted finite and infinite mixture sampling. The convoluted cases are exactly the same, but changing the equations with the modifications described in section S3.4. In addition to this, it is necessary to add a step in the sampling loop to sample new parameters from the already fitted noise distribution $\{\boldsymbol{\phi}_j^{\xi}\}_j$.

#### S6 Efficiency assessment

So far, the efficiency of deconvolution methods has been assessed using quantifiers applicable to point estimates, which are the ones proposed in the literature to date. To compare the efficiency of our model with previous methods that do not obey the positivity nor the normalization conditions, we can use the Mean Integrated Squared Estimation (MISE) measure:

$$\text{MISE} = \int_{-\infty}^{\infty} (p_{\text{true}}(\boldsymbol{x}) - p_{\text{est}}(\boldsymbol{x}))^2 dx, \tag{S101}$$

where $p_{\text{true}}$ is the real target distribution and $p_{\text{est}}$ is the point estimate of the deconvolved distribution. This is the traditional measure of convergence, but it is hard to interpret as it only has a lower bound. To address this issue, we introduce the mean integrated overlap (MIO):

$$\text{MIO} = 1 - \frac{1}{2}\int_{-\infty}^{\infty} |p_{true}(\boldsymbol{x}) - p_{est}(\boldsymbol{x})| dx \tag{S102}$$

This measure has the property that it is bounded in the interval $[0, 1]$ if the true and the estimated distributions are normalized, with 0 corresponding to the case of no overlap between distributions, and 1 to the case of complete overlap. Values below zero can be obtained if the estimated distribution does not follow the positivity requirement nor the normalization condition, as it is the case in FFT-based deconvolutions.

#### Supplementary figures

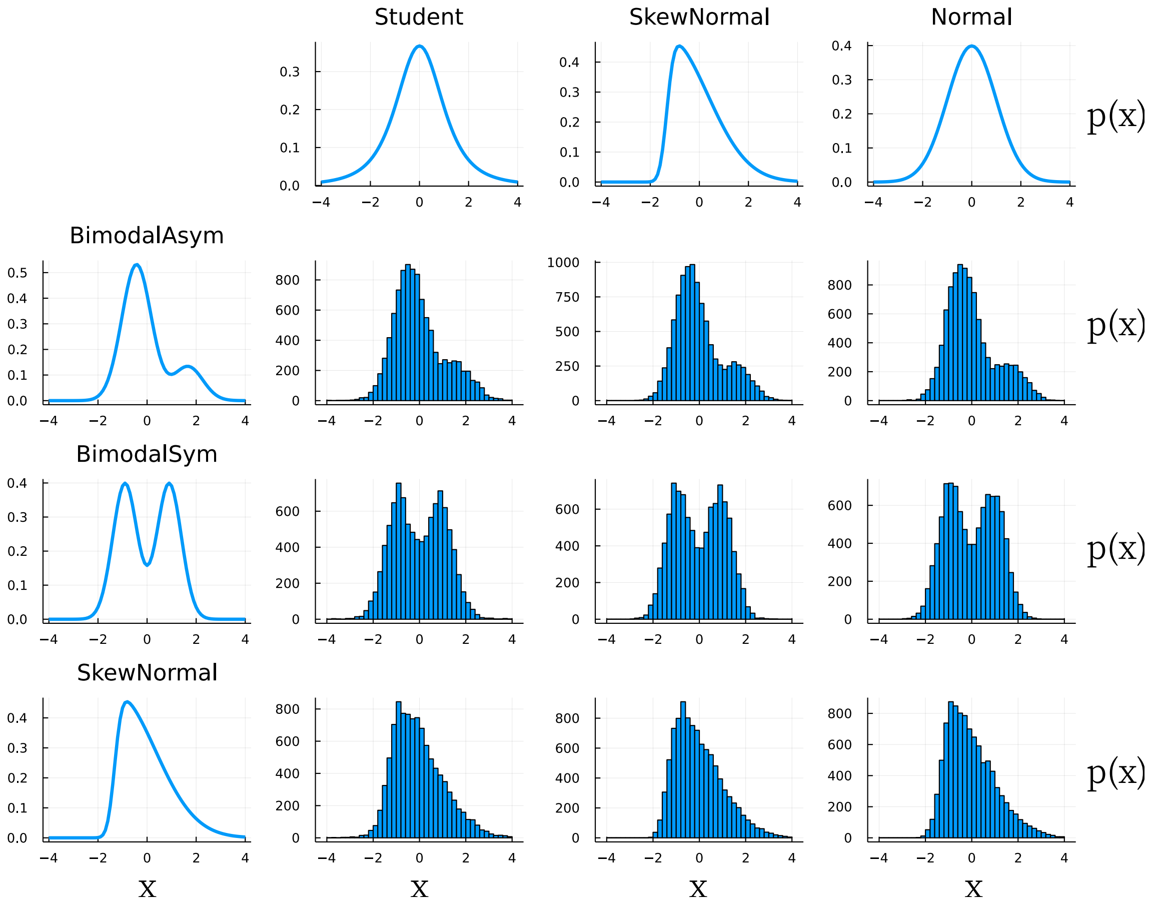

Figure S1: Synthetic target distributions (left) and noise distributions (top) and the resulting convolutions for a SNR=2.

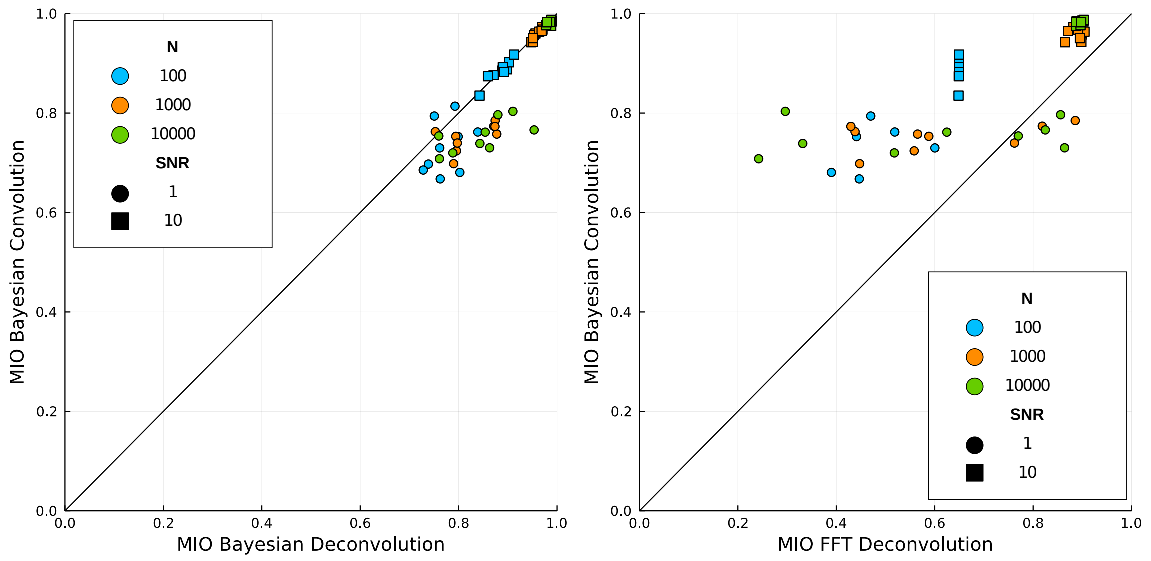

Figure S2: Comparison between the deconvolved and ground-truth target distributions as expressed by the Mean Integrated Overlap (MIO) for the Bayesian (left) and FFT (right) methods (x axis), with the results of a null Bayesian model fitting directly to the convolved data (y axis), ignoring the noise.

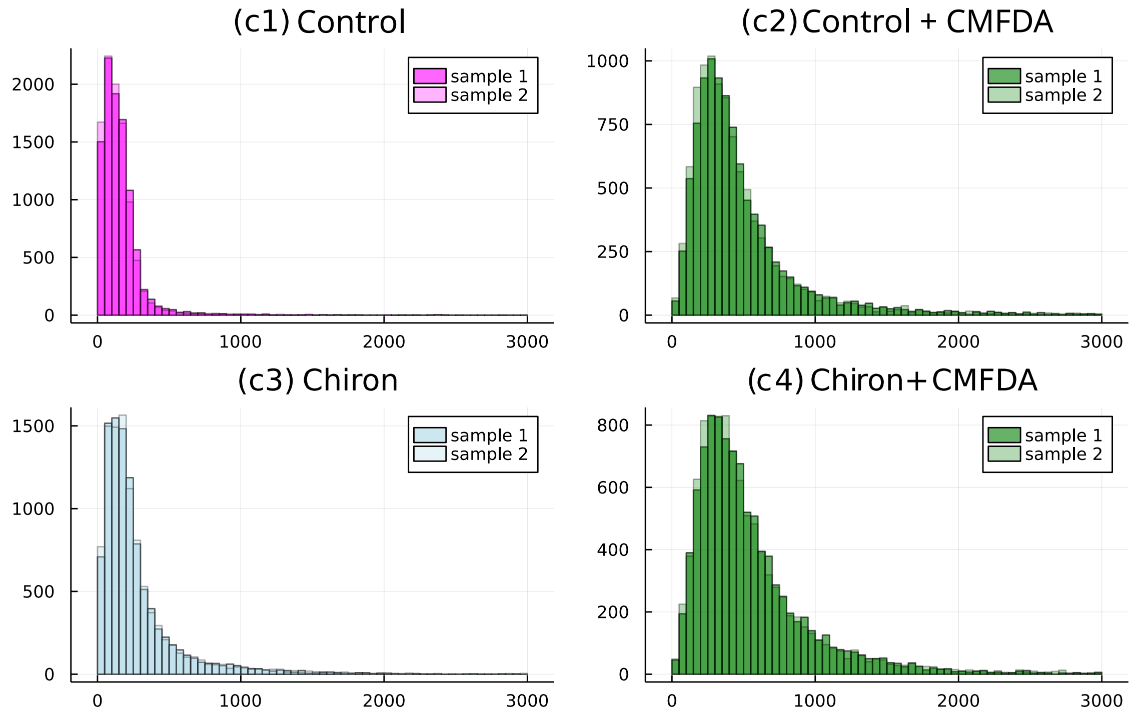

Figure S3: Flow cytometry distributions obtained in the four experimental conditions discussed in Sec. III.B of the main text.

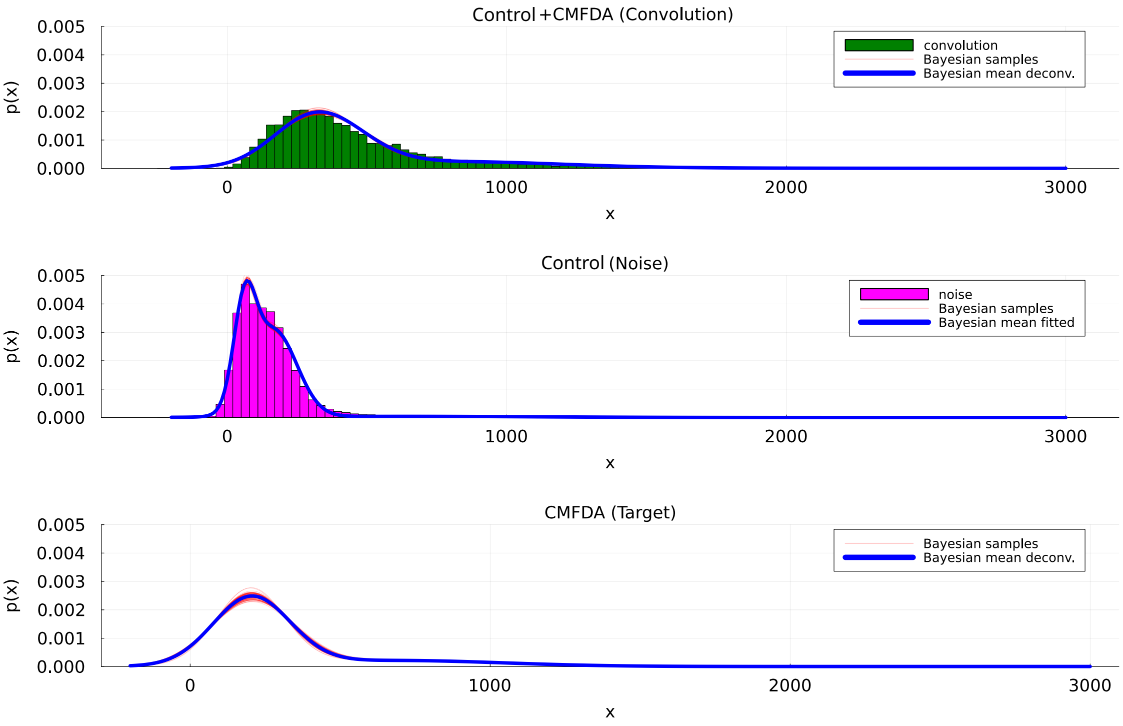

Figure S4: Flow cytometry distributions corresponding to the data conditions c2 (top panel in green) and c1 (middle panel in magenta) discussed in Sec. III.B. Overlayed on the distributions we show realizations of the Bayesian sampling process (red lines) for the three distributions (noise, convolution and target) obtained during the fitting.

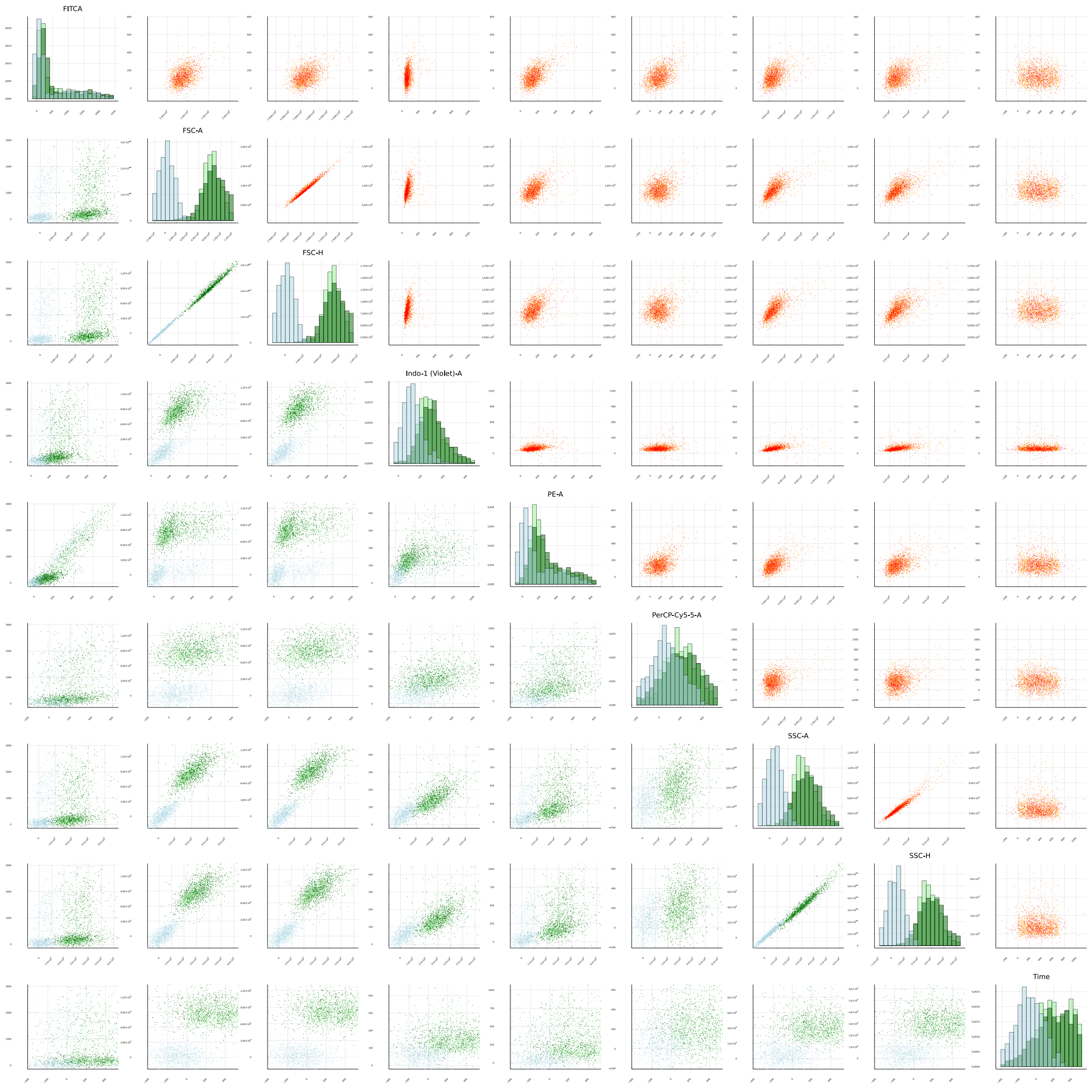

Figure S5: Scatter plot sections of the multichannel dataset. The top right triangle shows the autofluorescence distribution (orange) and samples from the fitting process (red). In the low left triangle we show the convolution data (light green) and samples from fitted convolution (dark green) and the deconvolution (light blue). In the diagonal, we show the 1D distributions of the convolved and deconvolved results.
